# ILC2s govern imprinting of alveolar macrophage-mediated immunity upon secondary helminth infection in the lung

**DOI:** 10.64898/2025.12.22.695921

**Authors:** Jonathan Pollock, Jhanvi H. Patel, Lisa-Marie Graf, Andreas Ruhl, Daniel Radtke, David Voehringer

## Abstract

Hookworm parasites migrate through the pulmonary system as part of their lifecycle, causing significant tissue damage that activates tissue-resident cell populations such as ILC2s. These cells drive type 2 immune responses critical for wound healing and the development of protective immunity. Macrophages, particularly monocyte-derived macrophages that seed the lung after infection, are central effectors of these responses and may be distinct from the tissue-resident populations they replace. However, the cellular and molecular factors governing the attrition of tissue-resident (TR) macrophages and their replacement remain poorly understood. We find that ILC2s and IL-4/IL-13-producing CD4^+^ T cells bi-directionally interact to regulate alveolar macrophage (AM) populations during infection. In the absence of these cells, type 2 immune responses are muted, leading to impaired anti-parasitic immunity in the lung during re-infection. This correlates with diminished expansion of type 2-polarized monocyte-derived alveolar macrophages (Mo-AMs), which are metabolically and transcriptionally distinct from TR-AMs. While Mo-AMs highly upregulated expression of Arg1, this was not essential for development of protective immunity in the lung. Concomitant with a more glycolytic metabolism, ILC2-induced Mo-AMs selectively upregulate arachidonate 15-lipoxygenase (Alox15), which promotes the capacity of macrophages to attack parasites in vitro. These results identify ILC2s as critical regulators of alveolar macrophage dynamics and important mediators of their function during helminth infection of the lung.

## Introduction

The human hookworms *Necator americanus* and *Ancylostoma duodenale* infect several hundred million people in countries with appropriate climatic conditions and poor hygiene standards. Larval stages of these parasitic worms penetrate the skin and migrate through the pulmonary system as part of their lifecycle^1^. Unlike other common helminthiases, the intensity of hookworm infections does not peak in childhood and people may become infected repeatedly^1^, which may lead to persistently altered lung function^2,3^. In laboratory models of hookworm infection, such as with the murine nematode *Nippostrongylus brasiliensis* (Nb), larval migration through the lungs leads to emphysematous lung pathology and chronic airway hyperresponsiveness, which is associated with accumulation of alternatively activated macrophages and caused by IL-4Rα-mediated activation of various cell types^4–6^ . The induction of these and other type 2 immune response in the lung is initially governed by IL-17A-elicited recruitment of neutrophils^7,8^, but is then driven by alarmin-activated type 2 cytokine producing innate lymphoid type 2 (ILC2s) and Th2 cells^9,10^. In the absence of these cells, muted pulmonary type 2 immune responses develop^5,11^ , which is associated with impaired recall immunity during secondary infection^12,13^. Thus, the induction of pulmonary type 2 immunity facilitates protection to subsequent infections by directly driving parasite killing in the lung, a phenomenon largely absent during primary infection^14^. Anti-parasitic immunity is thought to be principally driven by macrophages as macrophage-depleted mice exhibit enhanced susceptibility whereas macrophage-reconstituted mice exhibit enhanced larval killing^9,14^, though other cells such as eosinophils may also contribute^15^. These findings are supported by *in vitro* studies demonstrating that macrophages from Nb-infected mice are able to efficiently adhere to - and restrict viability of - Nb larvae^14,16^. While the precise mechanisms underlying macrophage killing of larvae in the lung remain elusive, their ability to restrict viability of larvae *in vitro* is thought to be driven by Arginase 1-mediated depletion of the essential amino acid arginine^16^. *Arg1* is differentially induced in lung macrophage subsets during Nb infection^16,17^, which has led to the hypothesis that macrophage populations do not uniformly contribute to pulmonary immunity in the lung, akin to the functional specialization and anatomical organization of different lung macrophages during homeostasis^18^.Under steady-state conditions, the major populations of macrophages in the lung, tissue-resident alveolar macrophages (TR-AMs) and tissue-resident interstitial macrophages (TR-IM) are derived from early yolk sac progenitors and fetal liver monocytes^19,20^ and retain the capacity to self-renew ^21^. As their names imply these macrophages inhabit distinct tissue niches within the lung with alveolar macrophages (AMs) residing in - or moving between - the alveoli^22,23^ and interstitial macrophages (IMs) residing in interstitial spaces, often in close association with blood vessels^24^ or nerve fibers^25^. When homeostasis of the lung becomes perturbed, such as during infection^26^, acute lung injury or allergic inflammation^27^, populations of tissue-resident macrophages can significantly diminish in number, as cell death processes outpace proliferation^3,28^. To maintain and restore lung function, lung macrophage populations are replenished by recruited monocytes which infiltrate pulmonary niches and differentiate into interstitial (termed Mo-IM) and alveolar macrophage (Mo-AM) populations^29^, often outcompeting extant tissue resident populations in the process^30^. The phenotype of these monocyte-derived macrophage populations is initially substantially distinct from tissue-resident populations derived during embryogenesis or early development, but becomes more similar over time^16^. In the context of an active immune or wound-healing response in the lung however, this generates a temporal window in which the responses of lung macrophage populations to microenvironmental factors, such as cytokines^31^, are notably altered. This can have either detrimental effects - exemplified by monocyte-derived macrophage driven lung fibrosis^32^ and infection-associated pathology^33^ - or beneficial consequences, as highlighted by studies showing enhanced anti-viral^34^, anti-bacterial^26^ or anti-tumor^35^ immunity in the lung following reshaping of the lung macrophage pool. Such effects also occur during pulmonary Nb infection^16,34^, where migration of larvae through lung tissue drives the emergence of *Arg1* expressing monocyte-derived alveolar macrophages, which preferentially aid in trapping of invading larvae ^16^.While recruited Th2 cells are known to be important in the development of protective monocyte-derived pleural macrophage populations during *L .sigmodontis* infection^31^ and mediate conversion of alveolar macrophages in various settings of allergic airway inflammation^36^, their precise role in restructuring alveolar macrophage populations during Nb infection remains ill- defined. Further, it remains elusive to what extent ILC2s influence this process, with recent studies showing that these cells have non-redundant functions^10,11^ and instead cooperate with Th2 cells to mediate type 2 immunity during Nb infection^9,37^. To gain insight into this, we study here the impact of ILC2 deficiency as well as conditional knockout of IL-4/IL-13 production in either KLRG1^+^ or CD4^+^ cells on priming pulmonary immunity to Nb. Our observations suggest that ILC2s, partially via their own production of type 2 cytokines, interact with Th2 cells in a bidirectional manner to shape alveolar macrophage populations. In the absence of both ILC2s as well as Th2 cells, alveolar macrophages populations are less heterogenous, maintain a phenotype consistent with TR-AMs and do not upregulate expression of *Alox15* - a regulator of anti-parasitic immunity ^38^.

## Results

### ILC2 deficient mice exhibit impaired pulmonary recall immunity to *N. brasiliensis* infection

ILC2s play established roles in regulating recall immunity during *N. brasiliensis* (Nb) infection in the small intestine^13^. However, whether the absence of ILC2s also functionally impacts on recall responses in the lung – an important site for priming of immunity ^14,39^- remains unclear. To investigate this further, we studied worm burdens in the lungs of Tie2^Cre^Rorα^fl/fl^ mice, which lack ILC2s^40^, and littermate controls two days after primary or secondary infection (40 days post primary infection). While worm burdens during primary infection were similar, Tie2^Cre^Rorα^fl/fl^ mice had significantly higher numbers of viable larvae in their lungs following secondary infection (Figure 1A). Consistent with this, Tie2^Cre^Rorα^fl/fl^ mice exhibited blunted type 2 immune responses, including lower mucus production (Figure 1B), fewer non-epithelial Arginase 1 and RELM-α producing cells (Figure 1D) and lower levels of the type 2 cytokines IL-4, IL-5, IL-9, and IL-13 (Figure 1E). In contrast, the expression of barrier response genes was only modestly affected (Figure 1C). In summary, these data suggest that ILC2s are important for promoting pulmonary recall responses during secondary Nb infection and impact on diverse type 2 immunity-associated effector functions.

**Figure 1.**
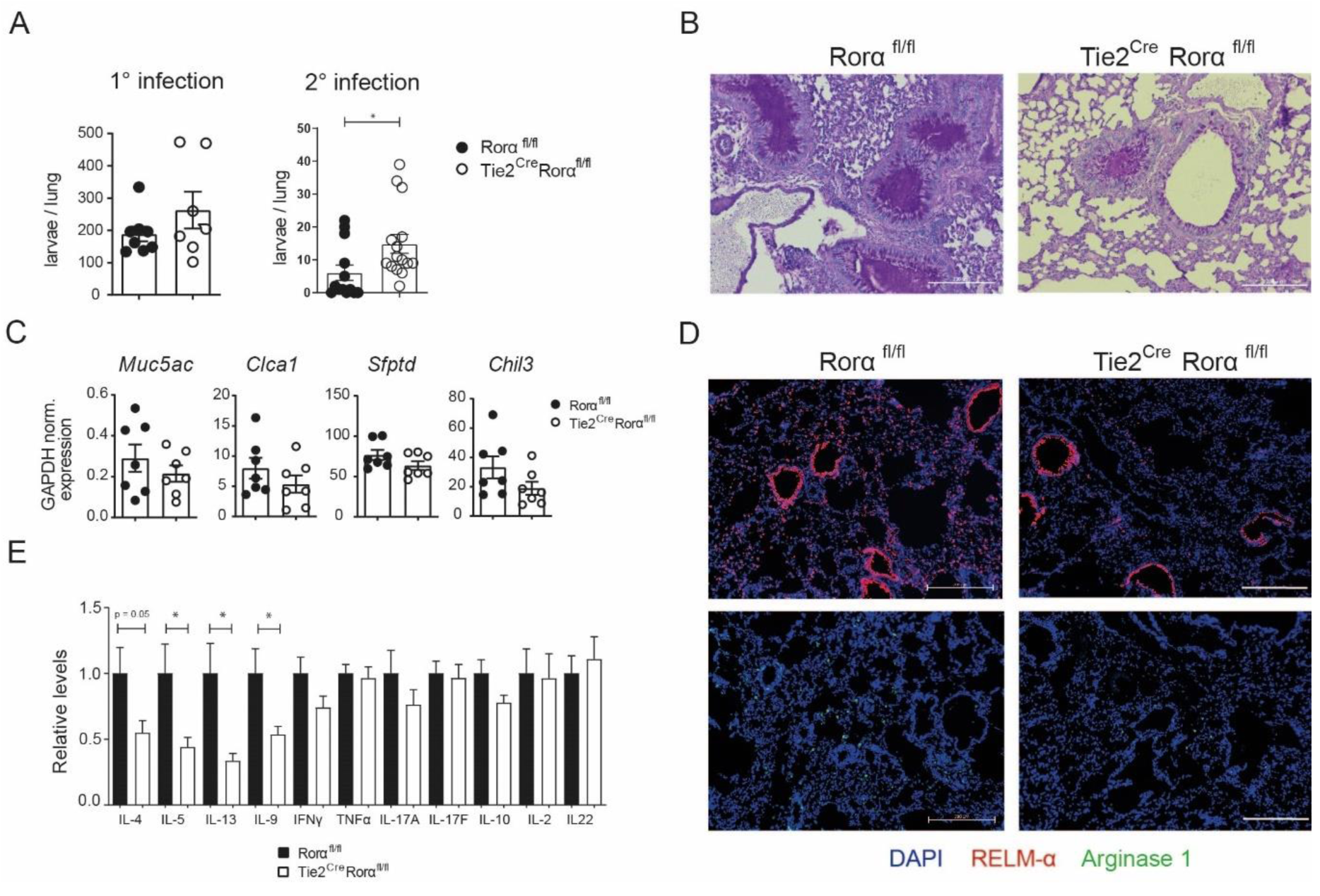
ILC2-deficient mice exhibit impaired pulmonary recall immunity to *N. brasiliensis* infection. Tie2^Cre^ Rorα^fl/fl^ and littermate control mice were infected with 500 L3 *N. brasiliensis* (Nb) larvae s.c and analysed at day 2 post primary (1°) or secondary (2°) infection (40 days post 1° infection). **A**) Number of larvae in the lungs of indicated mice. **B**) PAS staining of lung sections from indicated mice after 2° infection. Scale bars: 200 µm. **C**) qRT-PCR analysis of genes associated with mucosal type 2 immunity in total lung tissue of indicated mice after 2° infection. **D**) Immunofluorescent staining of RELM-α and Arginase 1 in cryosections of lung tissue from indicated mice after 2° infection. Scale bars: 200 µm. **E**) Cytometric bead array analysis of cytokine levels in lung tissue homogenates of indicated mice after 2° infection. Data in A and C are pooled from 2-4 independent experiments (n = 2-4 mice per group). Data in E are pooled from 4 independent experiments (n= 3-5 mice per group). Data in B and D are from one experiment, representative of two independent experiments. After testing for normality, unpaired Student’s t-tests were used for statistical analysis (*, p < 0.05). Bar graphs show the mean ± SEM.

### Absence of ILC2s impairs expansion of type 2 effector Siglec-F^lo^ alveolar macrophages and blunts eosinophilia

To further understand the immunologic consequences of ILC2 deficiency during secondary Nb infection of the lung we analysed several myeloid and lymphoid populations previously linked to pulmonary recall immunity^9,16,41^. As expected, Tie2^Cre^Rorα^fl/fl^ mice had vastly diminished relative and absolute levels of resident ILC2s compared to control mice (Figure 2A, Supplementary Figure 1 for gating strategies). Additionally, while levels of CD4^+^ and CD8^+^ T cells were similar (Supplementary Figure 2A-B), the relative and absolute levels of CD4^+^ST2^+^ (Th2) cells were also reduced in Tie2^Cre^Rorα^fl/fl^ mice (Figure 2B). Within the myeloid compartment we observed strong differences with respect to the relative levels and numbers (Supplementary Figure 2C-D) of alveolar macrophages (AMs) and eosinophils (Figure 2C) in Tie2^Cre^Rorα^fl/fl^ mice compared to controls. We also observed a consistent, but not significant, reduction in immature (CD11b^+^ SiglecF^-^Ly6G^-^ Ly6C^+^ MHCII^+^) macrophages in Tie2^Cre^Rorα^fl/fl^ mice, whereas other myeloid subsets were similar between groups (Figure 2D). Supporting an association of ILC2s with AMs, we found a significant negative correlation between relative levels of ILC2s and AMs in the lungs of both Tie2^Cre^Rorα^fl/fl^ and control mice (Figure 2E).

**Figure 2.**
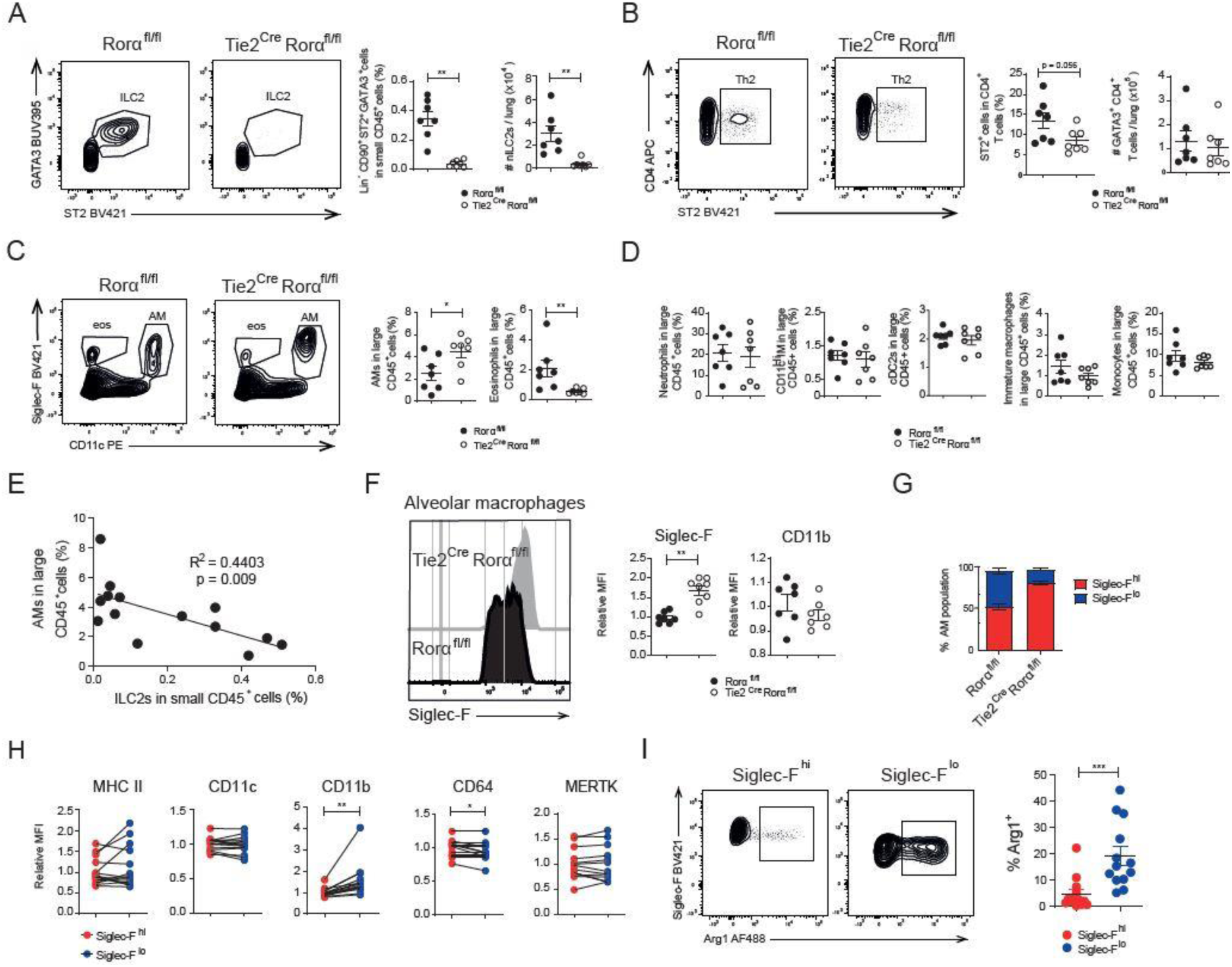
Absence of ILC2s impairs expansion of type 2 effector Siglec-F^Lo^ alveolar macrophages and blunts eosinophilia. Lungs of Tie2^Cre^Rorα^fl/fl^ and littermate control mice were analysed at day 2 post 2° infection with Nb larvae. **A**) Representative flow cytometry plots and quantification of ILC2s (CD45^+^Lin^-^ CD90^+^CD4^-^CD8^-^ ST2^+^GATA3^+^ cells) from indicated mice. **B**) Representative flow cytometry plots and quantification of Th2 cells (CD45^+^Lin^-^CD90^+^CD4^+^ST2^+^). **C**) Representative flow cytometry plots showing levels of eosinophils (eos, CD45^+^Siglec-F^+^CD11c^-^) and alveolar macrophages (AM, CD45^+^Siglec-F^+^CD11c^+^). **D**) Flow cytometric analyses of neutrophils (CD45^+^CD11b^+^Ly6G^+^), CD11c^hi^ inflammatory monocytes (IM, CD45^+^CD11b^+^CD11c^+^MHCII^+^CD64^+^), cDC2s (CD45^+^CD11b^+^CD11c^+^MHCII^+^CD64^-^), immature macrophages (CD45^+^CD11b^+^CD11c^-^MHCII^+^Ly6C^+^) and monocytes (CD45^+^CD11b^+^MHCII^-^Ly6C^+^). **E**) Linear regression analysis showing the correlation of alveolar macrophage and ILC2 levels in Tie2^Cre^Rorα^fl/fl^ and littermate control mice. **F**) Representative flow cytometry histograms showing expression levels of Siglec-F and CD11b on alveolar macrophages from indicated mice. **G**) Relative proportions of Siglec-F^hi^ and Siglec-F^lo^ alveolar macrophages (AM). **H**) Flow cytometric analysis of surface marker expression on SiglecF^Hi^ and SiglecF^Lo^ alveolar macrophages of Tie2^Cre^Rorα^fl/fl^ and control mice combined. **I**) Flow cytometric analysis of intracellular Arginase 1 expression in Siglec-F^hi^ and Siglec-F^lo^ alveolar macrophages. Data in A-I are pooled from two independent experiments (3-5 mice per group), representative of several experiments. Unpaired Student’s t-tests were used for statistical analysis in panels A-F and paired Student’s t-tests were used for data in H and I (*, p < 0.05; **, p < 0.01; ***, p < 0.001). Data in A-D, F and I include the mean ± SEM for each data set.

Interestingly, AMs from ILC2-deficient mice were also phenotypically distinct, showing much higher expression of Siglec-F but lower expression of CD11b (Figure 2F) compared to AMs from control mice, where Siglec-F^lo^ AMs also constituted a much greater fraction of the AM pool (Figure 2G). As expression of these markers are related to differences in AM development ^26^, we wondered whether Siglec-F^hi^ and Siglec-F^lo^ AMs also differed in their surface expression of other canonical macrophage markers.

While expression of MHC-II, CD11c and MERTK were similar between subsets, Siglec-F^lo^ cells were found to more highly express CD11b, and marginally less CD64 (Figure 2H). Interestingly, these defined AM subsets also markedly differed in terms of their intracellular expression of Arginase 1 (Figure 2I), suggesting differences in their capacity to polarize towards an alternatively activated phenotype, associated with anti-parasitic function^16^. Taken together, these data suggest ILC2s regulate AM heterogeneity during infection and are associated with the expansion of type 2-polarized Siglec-F^lo^ AMs – a signature consistent with nascent monocyte-derived AM (Mo-AMs).

### IL-4/IL-13 producing ILC2s and Th2 cells regulate AM populations during secondary infection

Differentiation and expansion of alternatively activated lung macrophages, crucial for resolution of lung pathology during Nb infection^42^, is known to depend on IL-4/IL-13 signalling^14,43^ . As Th2 cell-derived IL-4/IL-13 is also known to affect the development of monocyte-derived macrophages in the pleural cavity during helminth infection^31^, we hypothesized that absence of ILC2s impacts on AM populations by regulating pulmonary IL-4 and IL-13 levels. To study the relative contribution of IL-4/IL-13-producing ILC2s and Th2 cells in this regard, we studied the pulmonary immune responses to secondary infection of KLRG1^Cre^4-13^fl/fl^ and Cd4^Cre^4-13^fl/fl^ mice, which lack expression of IL-4 and IL-13 in either ILC2s or Th2 cells, respectively. KLRG1^Cre^4-13^fl/fl^ mice showed a trend of elevated worm burdens as compared to Cre^-^ negative littermates at day two after secondary infection (Figure 3A). While we observed similar levels of AMs as well as eosinophils in the lungs of KLRG1^Cre^4-13^fl/fl^ and littermate control mice (Figure 3B), AMs from KLRG1^Cre^4-13^fl/fl^ mice did show significantly altered expression of Siglec-F and CD11b (Figure 3C). Furthermore, while levels of resident ILC2s (Figure 3D) were unchanged in KLRG1^Cre^4-13^fl/fl^ mice we did observe a slight reduction in Th2 cells (Figure 3E). On the other hand, Cd4^Cre^4-13^fl/fl^ mice had significantly higher worm burdens at day 2 after secondary infection (Figure 3F) which coincided also with elevated levels of AMs (Figure 3G), similar to our observations in ILC2-deficient mice (Figures 1A, 2C). Consistent with our previous findings, AMs of Cd4^Cre^4-13^fl/fl^ mice also exhibited increased expression of Siglec-F and reduced expression of CD11b (Figure 3H). Finally, as expected, Th2 levels were also significantly reduced in Cd4^Cre^4-13^fl/fl^ mice (Figure 3J).

**Figure 3.**
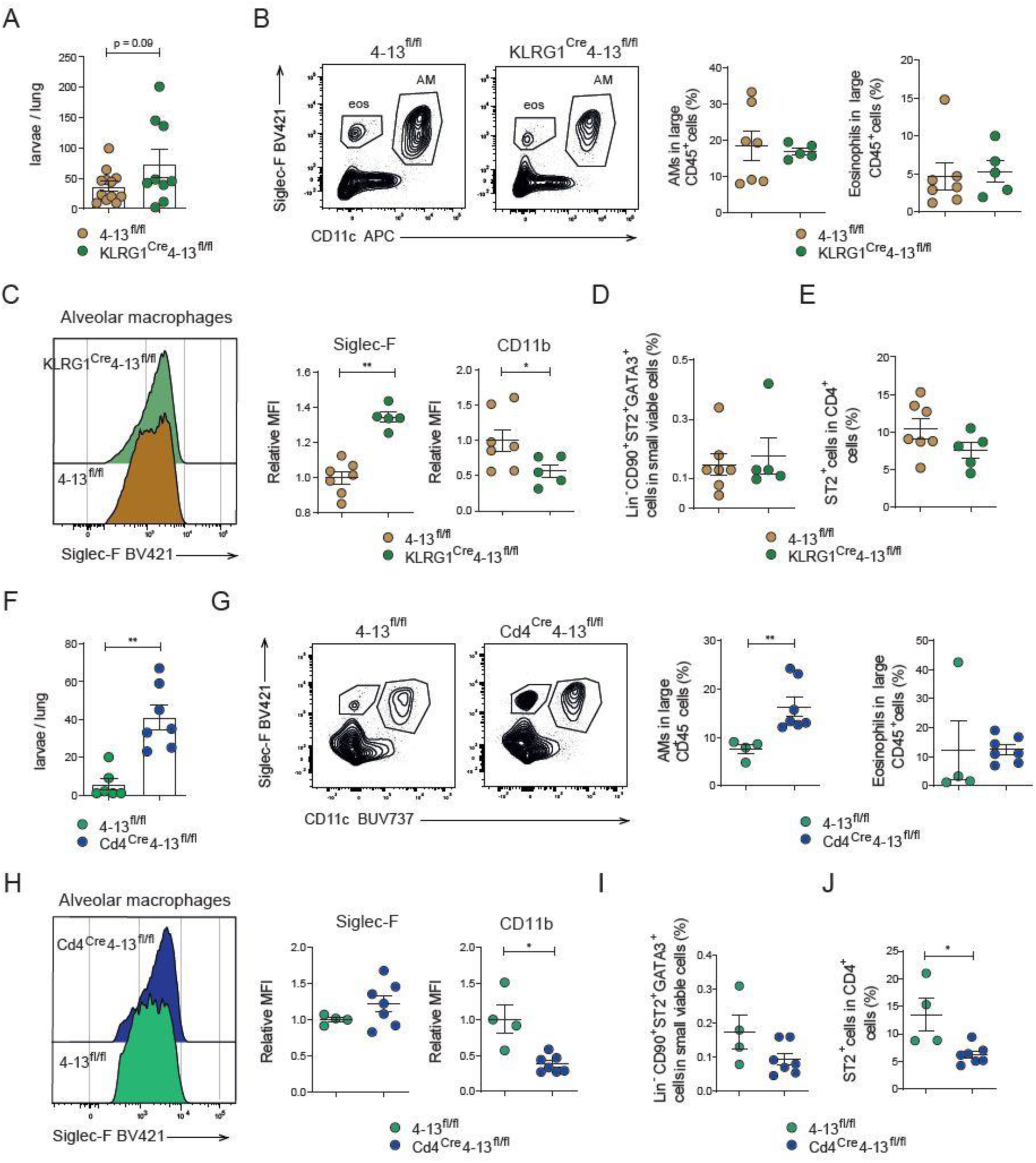
IL-4/IL-13 producing ILC2s and Th2 cells regulate alveolar macrophage populations during secondary infection. All data are from lungs of mice analysed at day 2 post 2° infection with Nb. **A**) Number of larvae in mice deficient in IL-4/IL-13 secretion from ILC2s (KLRG1^Cre^4-13^fl/fl^ mice) and littermate controls. **B**) Representative flow cytometry plots showing levels of eosinophils (eos, CD45^+^Siglec-F^+^CD11c^-^) and alveolar macrophages (AM, CD45^+^Siglec-F^+^CD11c^+^) in indicated mice. **C**) Representative flow cytometry histograms showing expression of Siglec-F and CD11b on alveolar macrophages from indicated mice. **D**) Representative flow cytometry plots showing levels of ILC2s (CD45^+^Lin^-^ CD90^+^CD4^-^CD8^-^ST2^+^GATA3^+^ cells) and E) Th2 cells (CD45^+^Lin^-^CD90^+^CD4^+^ST2^+^) in indicated mice. **F**) Number of larvae in mice deficient in IL-4/IL-13 secretion from CD4^+^ T cells (Cd4^Cre^4-13^fl/fl^ mice) and littermate controls. **G**) Representative flow cytometry plots showing levels of eosinophils and alveolar macrophages in indicated mice. **H**) Representative flow cytometry histograms showing expression of Siglec-F and CD11b on alveolar macrophages from indicated mice. **I**) Representative flow cytometry plots showing levels of ILC2s (and E) Th2 cells in indicated mice. Data in A are pooled from three independent experiments (2-5 mice per group); data in B-I pooled from two independent experiments (2-4 mice per group). Unpaired Student’s t-tests were used for statistical analysis (*, p < 0.05; **, p < 0.01). Data in A-I include the mean ± SEM.

Given the observed differences in AM populations seen in Cd4^Cre^4-13^fl/fl^ and Tie2^Cre^Rorα^fl/fl^ mice, and to a lesser extent in KLRG1^Cre^4-13^fl/fl^ mice, including fewer Arginase 1^+^ alveolar macrophages (Supplementary Figure 3A and B), we next sought to establish the role of Arginase 1 during recall immunity in the lung. While several previous studies have implicated Arginase 1 in regulating anti-parasitic responses of macrophages to Nb larvae *in vitro*^14,16^ and in the skin^44^ , it remains unclear to what extent endogenous Arginase 1 producing macrophages contribute to recall responses in the lung. To study this further, we analysed pulmonary immune responses during secondary infection of Tie2^Cre^Arg1^fl/fl^ mice, which broadly lack expression of Arginase 1 in hematopoietic cells including macrophages^45^. Unexpectedly, we observed no significant differences in pulmonary worm burdens between Tie2^Cre^Arg1^fl/fl^ and littermate control mice (Supplementary Figure 3C), despite vastly diminished Arginase 1 expression in lung tissue (Supplementary Figure 3D), as well as specifically in alveolar macrophages (Supplementary Figure 3E). Furthermore, flow cytometric analyses revealed similar levels of AMs and eosinophils (Supplementary Figure 3F), and similar proportions of Siglec-F^hi^ and Siglec-F^lo^ AMs (Supplementary Figure 3G) in Tie2^Cre^Arg1^fl/fl^ mice compared to littermate controls. This suggests that Arginase 1 expression marks macrophages with tissue-protective functions, but is not crucial for anti-parasitic immunity.

### scRNA-seq reveals transcriptional heterogeneity of alveolar macrophages and identifies immunologic and metabolic features associated with type 2 effector monocyte-derived macrophages

To gain insight into the transcriptional profiles regulating the distinct phenotypes of AM subpopulations, we performed scRNA sequencing of lung leukocytes at day 9 post primary infection. Previous studies have shown that infiltration of type 2 effector cells peaks around this time point^46^ and that these contribute towards wound healing as well as priming of secondary responses^9,39^. scRNA-seq resolved 23 separate clusters, comprising diverse lineages of myeloid, lymphoid and select stromal populations (Figure 4A). Consistent with our flow cytometric analyses, AMs comprised a greater fraction of cells in Tie2^Cre^Rorα^fl/fl^ and Cd4^Cre^4-13^fl/fl^ mice, whereas eosinophils were relatively less abundant in these mice relative to control mice (Supplementary Figure 4A and B). Sub-clustering analyses of cells with an AM/AM-like transcriptional profile resolved 5 different subpopulations, with distinct expression of canonical tissue-residency-associated (TR-AM) as well as monocyte-associated (Mo-AM) markers (Figure 4B-D).

**Figure 4.**
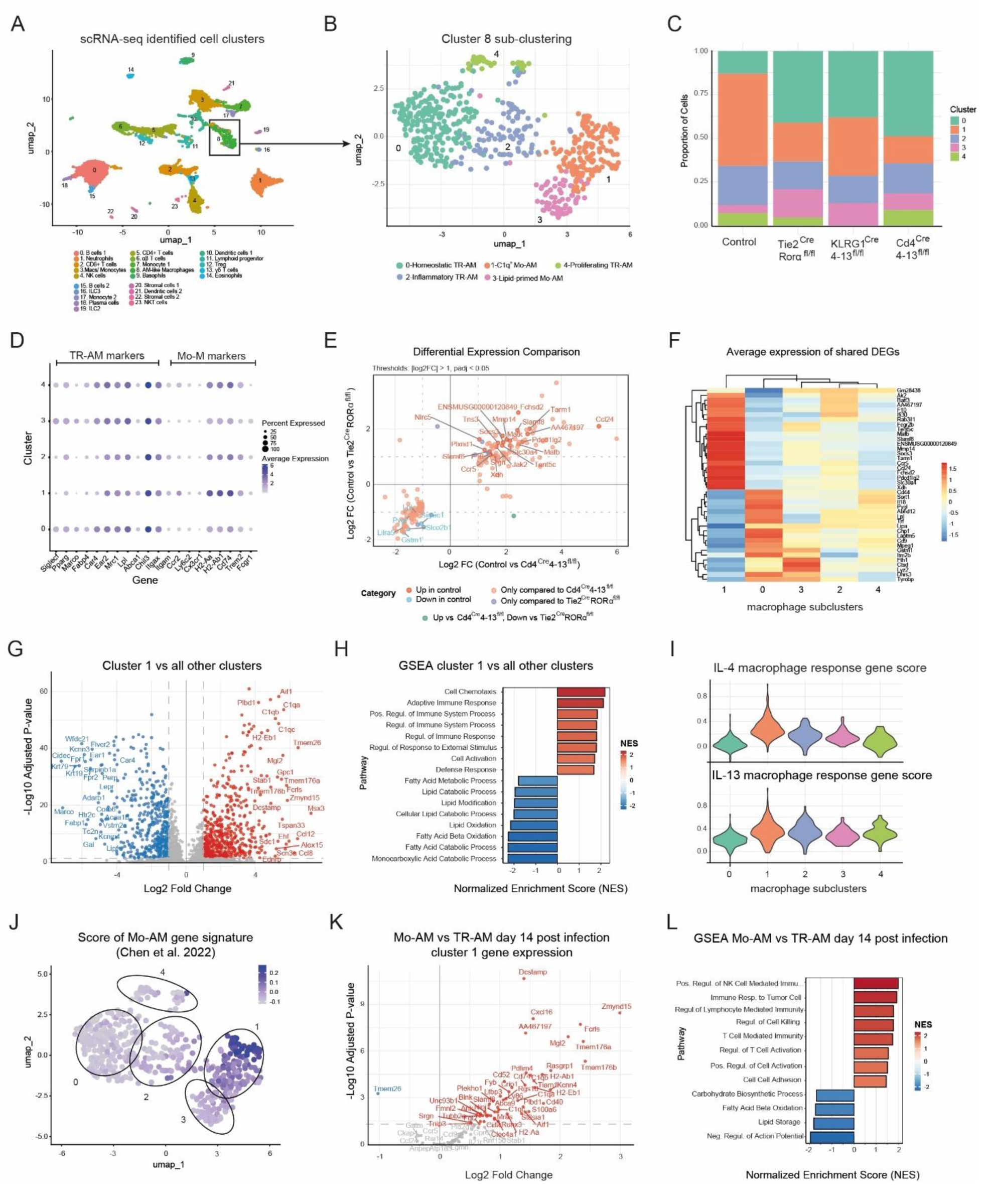
scRNA-seq reveals transcriptional heterogeneity of alveolar macrophages and identifies immunologic and metabolic features associated with type 2 effector monocyte-derived macrophages. Tie2^Cre^Rorα^fl/fl^, KLRG1^Cre^4-13^fl/fl^ , Cd4^Cre^4-13^fl/fl^ mice and littermate controls were infected with Nb and 9 days later lung single cell suspensions were subjected to scRNA-seq analysis. **A**) UMAP plot showing identified leukocyte and stromal cell populations. Cluster annotation was performed by inputting the top marker genes from each population into Immgen’s MyGeneset tool. **B**) Sub-clustering of AM-like cluster 8 from A). Cluster annotation was achieved by checking expression of canonical AM and Mo-AM marker genes, as shown in D, and with reference to published literature. AM: alveolar macrophages, Mo-AM: monocyte-derived AM, TR-AM: tissue-resident AM, IM: interstitial macrophages. **C**) Bar graphs show the relative proportion of each sub-cluster in cells from respective mouse groups. **D**) Dot plot shows frequency and average expression levels of markers associated with tissue resident alveolar macrophage (TR-AM) and monocyte-derived alveolar macrophages (Mo-AM) across respective clusters from B. **E**) Composite scatter plot shows differential gene expression between cells from control mice and cells from either Tie2^Cre^Rorα^fl/fl^ (y-axis comparison) or Cd4^Cre^4-13^fl/fl^ (x-axis comparison) mice. Only cells that met specified thresholds based on log2 fold change and adjusted p-value in at least one comparison are plotted and genes that are jointly up- or down-regulated in both comparisons are additionally labelled. **F**) Heatmap showing average expression of shared differentially expressed genes from E across sub-clusters from cells of all groups. **G**) Volcano plot shows a comparison of gene expression between cells of cluster 1 and cells of all other clusters. Significantly up- or down-regulated genes are coloured and top differentially expressed genes are additionally labelled. **H**) GSEA analysis shows statistically significant regulated pathways (GO BP database) based on expression profiles of genes from G. Red coloured bars represent pathways strongly associated with cluster 1 and blue coloured bars those strongly associated with other clusters. **I**) Violin plots showing expression levels of genes responsive to IL-4 and IL-13 treatment in macrophages of lymph nodes ^47^ across clusters and given as composite gene set scores. A higher score correlates with greater expression of genes regulated by either IL-4 or IL-13. **J**) Feature plot showing gene scores representing expression of Mo-AM signature genes (i.e. genes more highly expressed in Mo-AM vs TR-AM at day 14 post Nb infection), taken from a previously published study^16^ and overlayed on subclusters. **K**) Volcano plot analysis showing gene expression of the top 100 genes associated with cluster 1 from B between Mo-AM vs TR-AM . **L**) GSEA analysis showing statistically significant pathways (GO BP database) based on expression profiles of genes from J. Red coloured bars represent pathways strongly associated with Mo-AM and blue coloured bars those strongly associated with TR-AM. Data in A-I are from one scRNA-seq experiment (n = 2 mice per group). Data in J-L shows reanalysed data from one bulk RNA-seq experiment (n = 3 mice per group). For gene expression studies in E, G and K, Wilcoxon rank sum tests followed by Benjamini & Hochberg multiple-testing adjustment was used for statistical analyses.

Importantly, we observed marked differences in the relative proportions of these subpopulations in control mice relative to KLRG1^Cre^4-13^fl/fl^ and especially Tie2^Cre^Rorα^fl/fl^ and Cd4^Cre^4-13^fl/fl^ mice (Figure 4C). In line with this, differential gene expression analyses (all cells) of control versus Tie2^Cre^Rorα^fl/fl^ or control versus Cd4^Cre^4-13^fl/fl^ mice showed strong concordance (Figure 4E). Importantly, several of these concordant genes are known to be associated with type 2 immunity and were specifically expressed in certain sub clusters (Figure 4F), with many genes found to be more abundantly expressed in AMs from control mice being associated with cluster 1 (Figure 4G). Cells of this cluster were marked by high expression of the complement genes *C1qa*, *C1qb* and *C1qc*, exhibited low expression of *Siglecf*, but significant expression of *Itgam* (the gene encoding CD11b) (Figure 4D). To study these cells further we next contrasted their transcriptional profiles with those of all other subclusters and found that cells from cluster 1 highly expressed certain chemokines (*Ccl8*, *Cxcl16* and *Ccl12*), matrix metalloproteinases (*Mmp13* and *Mmp14*) and genes involved in antigen presentation (*H2-Aa*, *H2-Eb1* and *Cd74*) (Figure 4G). In contrast, cells from other clusters showed higher expression of classical TR-AM genes such as *Cidec*, *Krt79*, *Marco* and *Lepr* (Figure 4G). In agreement with this, functional enrichment analysis of these transcriptomic profiles demonstrated a strong functional association of cluster 1 with cell chemotaxis and regulation of immune responses (Figure 4H). Underscoring potential metabolic differences, cells from other clusters were enriched for pathways related to fatty acid metabolism (Figure 4H). Importantly, cells from cluster 1 also exhibited the highest expression (quantified as a composite gene set score) of genes associated with responses to IL-4 and IL-13 in macrophages (Figure 4I)^47^.

To confirm a monocytic origin of cells from cluster 1 we re-analysed a dataset from a previous study comparing gene expression profiles of Mo-AM vs TR-AM at day 14 post Nb infection^16^. From these data we generated a Mo-AM transcriptional signature (i.e. genes highly expressed in Mo-AM versus TR-AM) and found that cells from the AM subcluster 1 had highest expression of the Mo-AM signature (Figure 4J).

Complimentary to this, Mo-AMs also had significantly higher expression of many genes associated with cluster 1 (Figure 4K) and were positively enriched for pathways associated with immunoregulation (Figure 4L), but negatively enriched for fatty acid metabolic pathways. Taken together, these data suggest that loss of ILC2s or IL-4/IL-13 production from *Cd4* (and to a lesser extent *Klrg1*) expressing cells impairs the expansion of monocyte-derived AMs which have a strong type 2 immune signature and are specialized towards production of chemokines and the initiation of adaptive immune responses.

### Monocyte-derived effector alveolar macrophages are metabolically robust and mediate anti-parasitic responses via ALOX15

Tissue-resident (embryo-derived) AMs are hyporesponsive to type 2 cytokine stimulation due to their inability to efficiently use a range of metabolic substrates relative to monocyte-derived populations. To investigate the immunologic consequences associated with the impaired expansion of Mo-AM we sought next to characterize metabolic differences in AMs of ILC2-deficent Tie2^Cre^Rorα^fl/fl^ mice and littermate controls. In line with our scRNA-seq data, which showed a strong association of Mo-AM with expression of *C1qc* (Figure 4G), Tie2^Cre^Rorα^fl/fl^ mice exhibited diminished numbers of C1qc^+^ macrophages within- and surrounding alveoli (Figure 5A). To characterize metabolic differences between AMs of Tie2^Cre^Rorα^fl/fl^ and littermate control mice we isolated AMs two days after secondary infection and analyzed their incorporation of puromycin following treatment with metabolic inhibitors *ex vivo*^48^. Interestingly, in the absence of metabolic inhibition, AMs from Tie2^Cre^Rorα^fl/fl^ mice exhibited elevated incorporation of puromycin relative to AMs from control mice, but these differences diminished following treatment with either Oligomycin or 2-Deoxy-D-Glucose (Figure 5B). Further, while Siglec-F^hi^ and Siglec-F^lo^ AMs did not significantly differ in terms of their puromycin incorporation under control conditions, Siglec-F^lo^ AMs had higher puromycin incorporation during treatment with Oligomycin or 2-Deoxy-D-Glucose (Figure 5C), suggesting greater metabolic flexibility. Additionally, and in agreement with these data, we found that C1q^+^ Mo-AMs (cluster 1 in Figure 4B) showed lower expression of fatty acid oxidation genes, but higher expression of glycolytic genes (Supplementary Figure 5A-B) relative to cells with a TR-AM phenotype.

**Figure 5.**
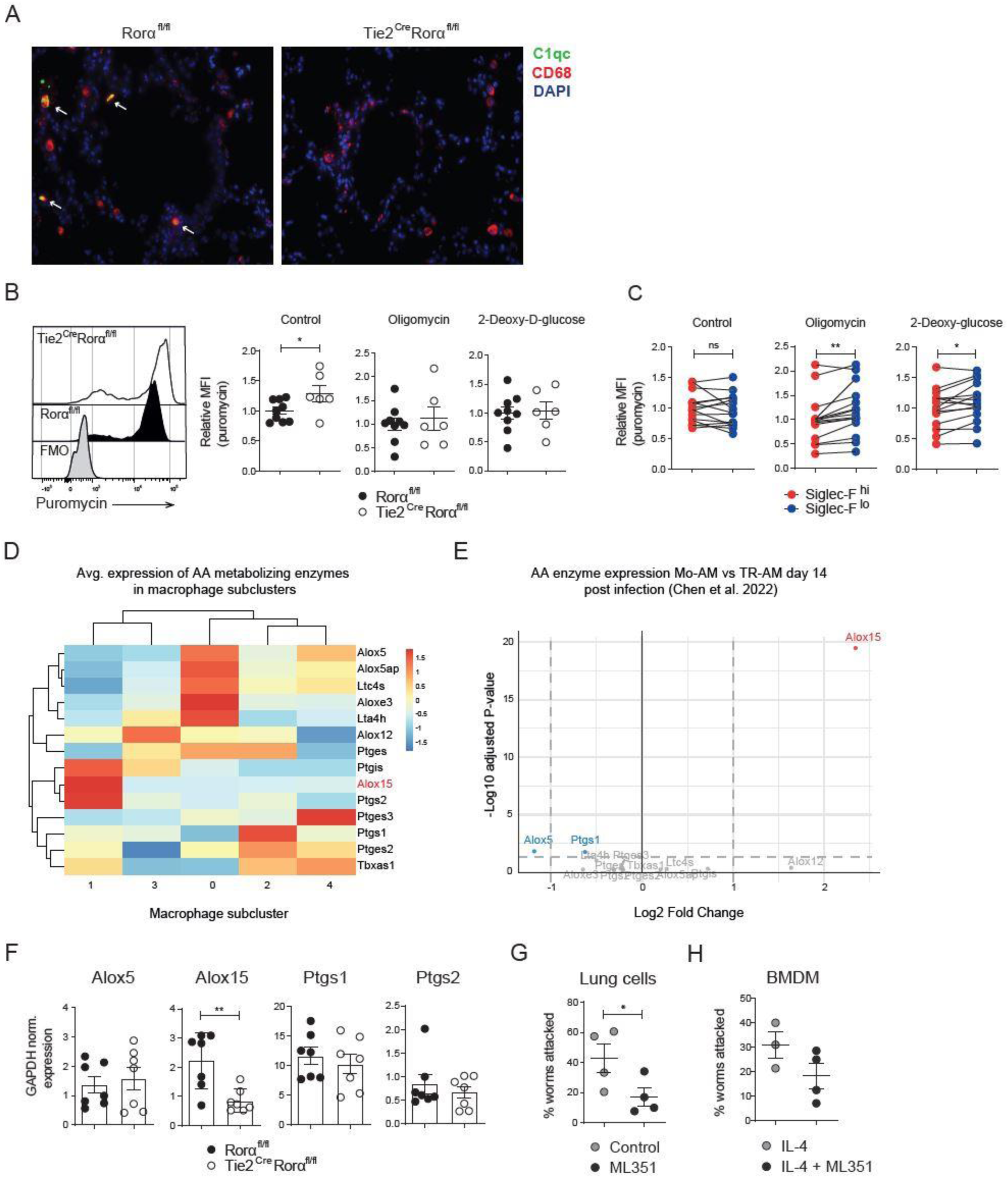
Monocyte-derived effector alveolar macrophages are metabolically robust and mediate anti-parasitic responses via ALOX15. **A**) Immunofluorescent staining of C1qc and CD68 (a pan macrophage marker) in cryosections of lung tissue from indicated mice after 2° infection. **B**) Representative flow cytometry histograms showing incorporation of puromycin in alveolar macrophages (AMs) of indicated mice. AMs were either treated with Oligomycin, 2-Deoxy-D-Glucose or vehicle control *ex vivo* prior to puromycin staining. **C**) Flow cytometric analysis of puromycin incorporation in Siglec-F^Hi^ vs Siglec-F^Lo^ AMs of Tie2^Cre^Rorα^fl/fl^ and littermate control mice. Cells were treated as in C. **D**) Heatmap showing relative gene expression of arachidonic acid (AA) -metabolizing enzymes in respective macrophage subclusters from Figure 4B**. E**) Volcano plot showing expression of (AA)-metabolizing enzyme gene expression in bulk transcriptomes of Mo-AMs versus TR-AMs at day 14 post infection. Statistically significant genes are coloured and labelled. **F**) qRT-PCR analysis of select AA-metabolizing genes in whole lung tissue from Tie2^Cre^Rorα^fl/fl^ and littermate control mice two days after 2° infection. **G**) *In vitro* larval attachement assay of lung leukocytes isolated from mice after 2° infection. Cells were either treated with the ALOX15 inhibitor ML351 (50 µM) or DMSO during co-culture with exsheathed L3 larvae and the % worms attacked were quantified after 48 hours. **H**) BMDM were generated from naïve WT mice and either polarised with IL-4 or with IL-4 + ML351 (10 µM) for 24 hours prior to co-culture with exsheathed L3 larvae. The % worms attacked were then quantified after 48 hours. Data in A from one experiment, representative of two experiments. Data in D are from one scRNA-seq experiment (n = 2 mice per group). Data in E show re-analysed data from one published bulk RNA-seq experiment (n = 3 mice per group). Data in B, C and F are pooled from two independent experiments (n = 3-5 mice per group). Data in G and H from one independent experiment. For gene expression studies in E, Wilcoxon rank sum tests followed by Benjamini & Hochberg multiple-testing adjustment was used for statistical analyses. For data in B and F unpaired Student’s t-tests and for data in C, paired Student’s t-tests were performed for statistical analysis. Data in B, C, F, G and H include the mean ± SEM. *, p < 0.05; **, p < 0.01.

Recent studies have highlighted how macrophage effector functions and their polarization state are regulated by broad metabolic changes^49,50^ and the activity of arachidonic acid metabolizing enzymes in particular ^38^. Gene expression analysis of the major arachidonic acid metabolizing enzymes in AM subclusters (Figure 4B) revealed significant differences, with cells from clusters with a TR-AM signature expressing higher levels of *Alox5* and *Ltc4s* and those with a Mo-AM signature, especially cells from cluster 1, expressing higher levels of *Alox15* (Figure 5D). Further, *Alox15* was highly expressed in Mo-AM but not TR-AMs at day 14 post Nb infection (Figure 5E), suggesting potential for these differences to persist after the acute phase of lung inflammation. In line with this, qRT-PCR analyses of genes involved in arachidonic acid metabolism showed a selective deficiency of *Alox15* in whole lung tissue of Tie2^Cre^Rorα^fl/fl^ mice (Figure 5F). Finally, to investigate the functional importance of Alox15 in regulating anti-parasitic responses we studied how inhibition of this enzyme affects the capacity of macrophages to attack parasitic larvae *in vitro*. We found that treatment of lung leukocytes isolated from mice two days after secondary infection with the ALOX15 inhibitor ML351 ^51^ significantly reduced the proportions of attacked larvae (Figure 5G). Additionally, bone-marrow derived macrophages (BMDMs) which were treated with ML351 during polarization with IL-4^38^ were also found to be less capable at adhering to larvae (Figure 5H). Taken together, these data suggest that the loss of ILC2s impacts on the metabolic phenotype of AMs after infection and implicate Alox15 as a relevant regulator of anti-parasitic responses.

## Discussion

In this study we have discovered important cellular and molecular regulators shaping alveolar macrophage (AM) populations during Nb infection. In line with previous studies, our findings demonstrate an important role for Th2-derived IL-4/IL-13 in driving the emergence of monocyte-derived alveolar macrophage populations in the lung, which are known to more readily adopt an alternatively activated phenotype^16,50^. Extending these findings, we also find that lung-resident ILC2s contribute towards regulation of AMs during Nb infection, partially through their own production of IL-4/IL-13, which is in line with previous studies demonstrating ILC2s constitute a substantial fraction of IL-13^+^ cells during secondary infection of the lung^9^. Further, given our previous findings highlighting macrophage-intrinsic STAT6 activation as a critical feature regulating the ‘disappearance reaction’^52^ of alveolar macrophages in allergic airway inflammation^36^, it is likely that Th2- and ILC2-derived IL-4/IL-13 acts directly on alveolar macrophages to regulate their fate and function. In *L. sigmodontis* infection, IL-13 has a dominant role in regulating the number – and differentiation of serous-cavity macrophages from monocyte precursors, which are critical for anti-parasitic immunity^31^. Similarly, our results indicate that in the absence of ILC2 or Th2-derived IL-4/IL-13, monocyte-derived alveolar macrophage populations do not optimally establish in the lung. This is likely due to several concomitant effects these cytokines have on AMs. Notably, recent studies have indicated that these cytokines can, through a yet to be discovered mechanism, directly diminish AM populations^36^, thus providing an opportunity for recruited monocyte-derived macrophages, whose proliferation is supported by IL-4Rα signaling^53^, to replenish the niche. Other studies also indicate that IL-4Rα signaling supports survival of monocytes in the periphery^54^. However, as we did not detect differences in the levels of monocytes between ILC2-deficient and conditional IL-4/IL-13 deficient mice and littermate controls, this is unlikely to be the major factor driving discrepant emergence of Mo-AMs. Instead, and in line with our observations showing heightened proliferation of AMs in ILC2-deficient mice, we hypothesize that TR-AMs either outcompete incoming recruited macrophages or otherwise restrict their ability to seed the lung when IL-4/IL-13 levels are diminished. Both ILC2s and Th2 cells seem to be necessary for optimal production of type 2 cytokines and bidirectionally support each other, as evidenced by lower Th2 cells in Tie2^Cre^Rorα^fl/fl^ and lower ILC2s in Cd4^Cre^4-13^fl/fl^ mice. This is in keeping with previous studies showing impaired ILC2 expansion and activation following depletion of CD4^+^ T-cells prior to infection^9^, and lends further support to these cells cooperating in priming development of protective immunity in the lung^39,41^.

Macrophage heterogeneity is emerging as an important feature affecting immune responses to infectious agents in the lung. A dominant driver of this heterogeneity, alongside microenvironmental cues^55,56^, is macrophage ontogeny^20^. A common phenomenon following acute lung infection, especially during severe infection, is the re-seeding of the alveolar macrophage-compartment with monocyte-derived cells. While the transcriptional profiles of both Mo-AM as well as TR-AM are largely specified by the microenvironment^57^ some imprinting as a result of their differential origin remains^58^ .In Nb infection Mo-AM and TR-AM have a markedly distinct phenotype shortly after larval migration through the lung (day 7), but become more similar by day 14 post infection^16^. In this study, we have tracked these different populations through their differential expression of the canonical AM marker Siglec-F^59^ and show that differences between subsets persist until secondary infection. In particular, we found that Mo-AM and TR-AM exhibited differences in their metabolism, consistent with previous studies looking at metabolism of macrophage subsets in the small intestine during Nb infection^60^.

Importantly, these metabolic differences, including potential differences in glycolytic metabolism, are known to alter responsivity of AMs to type 2 cytokines such as IL-4^56^. This may partially explain why Mo-AM exhibit a greater potential to polarize towards an alternatively activated (M2) state and thereby drive type 2 immune responses^33^. A classical marker of M2 macrophages is the metabolic enzyme Arginase 1, which is believed to kill parasitic larvae by directly depleting the essential amino acid arginine^16^. In contrast to previous studies however, we were unable to show a role for this enzyme in regulating pulmonary immunity to Nb infection, which may suggest it is strongly upregulated in macrophages for other reasons – such as facilitating wound healing and promoting tissue repair. These findings are supported by a recent study, which showed that *Arg1* expression and the production of urea can de decoupled from macrophage anti-parasitic responses^38^. Instead, macrophage expression of the metabolic enzyme arachidonate 15-lipoxygenase , which catalyzes the peroxidation of polyunsaturated fatty acids (PUFAs) – a process that facilitates non-immunogenic apoptosis and production of lipid mediators with anti-inflammatory activity^61^-has emerged as an important driver of macrophage anti-parasitic responses^38^. Complimenting studies demonstrating a close link between Alox15 and IL-4 in human monocyte-derived macrophages^61^, we show that macrophages with an Mo-AM signature specifically upregulated Alox15 relative to other AM subsets and that blockade of this enzyme hindered the capacity of macrophages to attack larvae *in vitro*. While we do not exclude the possibility that Alox15 production in other cells, such as neutrophils^62^, also play relevant roles in regulating AM function, our data indicate an important function of alveolar macrophage-intrinsic Alox15 expression. Alox15 has pleiotropic effects in macrophages, including regulating PPAR-γ signaling^63^, glycolytic metabolism^38^ and production of immunoregulatory molecules such as chemokines^64^.Our data point to the upregulation of this enzyme as a potentially important feature of macrophages that have polarized towards an M2 state, which facilitates their recognition and binding of parasitic larvae. Other transcriptional features associated with M2 polarized Mo-AM in this study, including elevated expression of C1q molecules and high expression of genes such as *Aif1* and *Plbd1* (among others), require further inquiry. Previous studies have shown that C1q production is not important for immunity to Nb^15^ or for altering responses of AMs to type 2 cytokine stimulation^65^, suggesting a non-essential function. AIF1 is upregulated in macrophages in a variety of (sterile) inflammatory contexts^66^, whereas PLBD1 is emerging as a promising biomarker in cancer studies^67^. Both of these impact on macrophage metabolism, with AIF1 linked to enhancing glycolytic metabolism^66^ and PLBD1 involved in lipid metabolism ^67^As such, monocytes adopting a Mo-AM phenotype exhibit a mixed metabolic phenotype which may allow them to respond to shifting microenvironments with a greater degree of flexibility relative to TR-AM^58^, and thereby carry out specific functions. This may include the production of chemokines which stimulate recruitment of other immune cells, such as eosinophils^68^, to the lung.

Consistent with this, we found that *Ccl24* (eotaxin-2) expression was significantly lower in AMs from Tie2^Cre^Rorα^fl/fl^ and Cd4^Cre^4-13^fl/fl^ mice, which associated with difference in eosinophil recruitment. Functional enrichment analyses further showed that cells with a Mo-AM signature also were enriched for pathways involving adaptive immunity and immune cell activation, but also exhibited elevated expression of certain immunosuppressive molecules, including PD-L2. These findings illustrate the potential dual-nature of Mo-AMs in the lung following Nb infection: They can abet inflammatory processes through stimulating recruitment and activation of other immune cells, but also play important roles in wound healing through production of soluble mediators and surface expression of immunosuppressive molecules.

## Materials and methods

### Mice

Tie2^Cre^Rorα^fl/fl^ ^69^, CD4^Cre^4-13^fl/fl^ ^70^ and Klrg1^Cre^ ^71^ mice have been described. Klrg1^Cre^4-13^fl/fl^ mice were generated by inter-crossing. Tie2^Cre^Arg1^fl/fl/^ and Tie2^Cre^RORα^fl/fl^ mice were kindly provided by J. Mattner and S. Wirtz, respectively (both University Hospital Erlangen). C57BL/6 mice were purchased from Charles River, Germany. Mice were housed under specific pathogen-free (SPF) conditions and mice of both sexes were used for experiments at the age of 8 – 16 weeks. For individual experiments, groups were sex and age matched. All animal experiments were performed in accordance with the German animal protection law and the guidelines of the European Union (Directive 2010/63/EU) with the permission of the Government from Lower Franconia.

### Nippostrongylus brasiliensis infection

Mice were infected with 500 L3 larvae in 200 µl 0.9% NaCl subcutaneously (s.c.) in the scruff of the neck. In secondary infection experiments, mice were reinfected with 500 L3 larvae s.c. five weeks later. To determine pulmonary larval burdens, whole lungs of infected mice were then finely minced, wrapped in gauze and suspended in 15 ml reaction tubes containing 0.9% NaCl at 37°C for several hours to let larvae crawl out and settle at the bottom of the tube. They were then enumerated using a stereomicroscope.

### Periodic acid-Schiff staining of lung sections

After overnight fixation in 4% paraformaldehyde (PFA), lung tissue were embedded in paraffin, sectioned at a depth of 3 µm and transferred to glass slides. Sections were then dewaxed and rehydrated by two times incubation for 5 min in xylol followed by 3 min incubations in 100% ethanol, 80% ethanol, 50% ethanol and finally distilled water. Glass slides were then incubated for 7 min in 0.5% periodic acid and rinsed two times in distilled water. Next, slides were immersed for 15 min in Schiff’s reagent, followed by three washing steps in sulphite water for 2 min. After counterstaining with Mayer’s haematoxylin, glass slides were mounted with Entellan and sections were analysed on an Axio Vert.A1 microscope.

### Immunofluorescent staining of lung sections

After overnight fixation in 4% PFA, lung tissue samples were washed in PBS and then incubated overnight in 25% sucrose (in PBS) solution at 4°C. The tissue was dried on paper towels, frozen on dry ice in cryomolds filled with OCT tissue freezing medium and stored at -80°C. Sections were then cut on a CryoStar NX70 at a thickness of 7 µm, transferred to StarFrost adhesive microscope slides dried for 1 h and then frozen or used for staining.

Slides were then incubated in 0.1% Triton X-100 in PBS for 10 min to permeabilize tissues, washed in 1x PBS-T (1x PBS, 0.1% Tween-20 ) and covered with TNB buffer containing 1 µg of α-CD16/32 Fc block antibodies and 3% normal donkey serum for 2 h to block non-specific binding. After washing in PBS-T, slides were then covered in primary antibodies diluted in TNB buffer overnight at 4°C. The next day, three washing steps were performed and secondary antibodies (1:500) were added together with DAPI (1:1000) in TNB buffer for 1 h at room temperature. After a final wash in PBS-T, slides were covered in Fluoroshield and mounted with cover slips. Images were acquired on an Axio Vert.A1 running Zen Blue edition software. To create composite image overlays, the add-channel function was used on images of separate fluorescent channels. A threshold was then applied for each channel based on background signal intensity in fluorescence-minus-one (FMO) control slides.

### Flow cytometry

Prior to collection of lung tissue, the lung was flushed with ∼3 ml of cold PBS supplemented with 2 mM EDTA. Lung tissue samples were then collected in collection media (RPMI 1640 supplemented with 10% FCS, Pen/Strep and 1x GutaMax) and minced with scissors. Tissue samples were then digested with Liberase and DNAse I (both 1mg/ml) at 37°C for 25 minutes with shaking. The digested tissue was then meshed through a 100 µm strainer and washed with PBS containing 2% FCS. After centrifugation, red blood cells were lysed by incubating cell pellets in 1 ml of ACK lysis buffer for 3 min at room temperature. Lysis was then stopped through addition of 30 ml of ice cold PBS and the samples were centrifuged, filtered again through a 70 µm strainer and resuspended in FACS buffer (PBS, 2% FCS, 0.1%NaN3). An aliquot was then taken of each sample and cell counts were enumerated using a Cellometer Auto T4 cell counter. 1.5 – 2x10^6^ cells were then transferred to 96-well U-bottom plates and non-specific staining was blocked by incubating cells in 100 µl of FACS buffer with α-CD16/32 (1:100) antibodies. Cells were then surface-stained with primary antibody cocktails (see antibody tables for specific details) for 45 minutes on ice in the dark. In some experiments, cell suspensions were then fixed and permeabilized using the Foxp3 intracellular staining kit fixative for 20 minutes on ice. After fixation and permeabilization, cells were stained with intracellular antibodies diluted in Foxp3 wash buffer overnight at 4°C, washed and resuspended in FACS buffer.

For puromycin incorporation staining, single cell lung suspensions were generated as described above and 1x10^6^ cells were transferred to 96-well U-bottom plates and resuspended in media (RPMI 1640, 2% FCS, Pen/Strep, 1x GlutaMax) and left to equilibrate for 30 min at 37°C and 5% CO2. After equilibration, cells were spun down and resuspended in either control media (RPMI 1640, 2%FCS, Pen/Strep, 1x GlutaMax, 0.01% DMSO) or in control media with addition of either Oligomycin A (10 µM final concentration) or 2-Deoxy-D-glucose (100 mM final concentration), and incubated for 15 min at 37°C and 5% CO2. Puromycin (10 µg/ml) was then added to each well (except control wells) and cells were left to incubate for a further 30 min. Cells were then spun down and washed in ice cold PBS before being surface and intracellularly stained as described above. All cells were acquired on a BD LSR Fortessa flow cytometer and analyzed using FlowJo Version 10.6.2.

### RNA isolation and real-time quantitative PCR

Lung tissue sections were collected in RNAprotect tissue reagent and kept on ice until processing. To isolate RNA, lung tissue sections were then briefly washed in sterile PBS and transferred to 2 ml screw-cap microcentrifuge tubes with 1 ml of RLT buffer. Two tungsten carbide beads were then added to the tubes and tissue was homogenized using a bead ruptor. RNA was then isolated using the RNeasy Mini kit by following manufacturer instructions. 2 µg of RNA was then used to generate cDNA using the High-Capacity cDNA Reverse Transcription kit.

Quantitative real-time polymerase chain reaction (qRT-PCR) was performed with SYBR Select Master Mix, primers indicated in Supplementary Table 1, and the following program: denaturation for 10 min at 95°C followed by 40 cycles of PCR amplification (95°C, 30 sec; 60°C, 45 sec; 72°C, 60 sec) and a melting curve (0.05°C/sec, 60-95°C). *CT* values were determined within QuantStudio Real-Time PCR software and expression of target genes was normalized to expression of the housekeeping gene *Gapdh using* the formula 2^-^(ΔCT).

### Cytokine quantification in whole lung tissue

Whole lung tissue was transferred to 1ml of RIPA buffer (1% NP-40, 50 mM TRIS pH 7.4, 0.15 M NaCl, 1 mM EDTA pH 8.0, 0.25% deoxycholic acid) containing a protease inhibitor cocktail and homogenized using a bead ruptor. The supernatant was then transferred to new tubes and total protein was quantified using a Bicinchoninic acid (BCA) assay. Cytokine levels were quantified using a 12-plex LEGENDplex™ kit (Mouse T Helper Cytokine Panel Version 3) by following manufacturer’s instructions. The beads were then acquired on a BD LSR Fortessa flow cytometer and cytokine levels quanitifed with LEGENDplex analysis software.

### Larval attachment assay with lung cells or bone marrow-derived macrophages (BMDM)

Lung leukocytes isolated from mice 2 days after secondary infection were co-cultured with larvae as described previously ^38^. Briefly, single cell suspensions were generated as per the protocol used for flow cytometric analysis and 5 – 10^6^ cells from cell pellets were resuspended in 12 ml of culture media (RPMI 1640, 10% FCS, Pen/Strep, 1x GlutaMax) and plated in sterile petri dishes overnight. Non-adherent cells were removed and adherent cells were scraped off of plates in ice cold PBS. 2x10^5^ adherent cells (largely macrophages) were then transferred to flat bottomed 96-well plates and resuspended in 50 µl of culture medium. Then 50 exsheathed ^14^ L3 larvae, which had been washed at least 3 times and diluted in culture media, were added in 50 µl of culture media. Plates were then incubated for 48 hours and the proportion of larvae that were attacked (defined as having several cells attached) was determined using a stereomicroscope.

BMDM were generated as described previously ^72^. Briefly, the tibias and femurs from 8-week-old mice were isolated and the bone marrow flushed through a 70 um strainer in PBS + 2% FCS. After washing the cells were resuspended in macrophage media (DMEM high glucose media, 10% FCS, Pen/Strep, 1x GlutaMAX) supplemented with 50 ng/ml rmM-CSF at a density of 12x10^6^ cells per 20 ml, plated on sterile petri dishes and cultured for 7 days. After 3 days an additional 12 – 20 ml of macrophage media was added to respective plates. At day 6 of the culture, IL-4 (10 ng/ml) alone or in combination with the ALOX15 inhibitor ML351 (10 µM) was added to the plates and the BMDMs were incubated for another 24 hours. Finally, BMDMs were scraped off the plate and incubated with larvae as described above.

### Single-cell RNA sequencing of lung leukocytes

Lungs were harvested at day 9 post primary infection and single cell suspensions were generated as described before. Dead cells were removed using the Dead Cell Removal Kit. Samples from two mice for each group were stained with the viability markers Calcein and DRAQ7 and multiplexed with hashtag antibodies according to the BD Single-Cell Multiplexing Kit. A total of 60,000 cells (7,500 cells per mouse) was used to load the cartridge before addition of beads and induction of cell lysis as specified in the “Single-Cell Capture and cDNA Synthesis with BD Rhapsody Single-Cell Analysis System” protocol. Following reverse transcription, library preparation was performed as outlined in the “mRNA Whole Transcriptome Analysis (WTA)” and “Sample Tag Library Preparation” protocols. The quality of amplification products was then verified on the Agilent TapeStation 4200 system using the Agilent High Sensitivity D5000 ScreenTape Assay. Libraries were sequenced on a NovaSeq 6000 instrument in the Next Generation Sequencing Core Unit of the Institute of Human Genetics of the University Hospital Erlangen, Germany.

### Single cell data processing

Data was processed with the BD Rhapsody WTA Analysis Pipeline version 1.12 and loaded into R for further analysis. Cell exclusion criteria were as follows: cells had to have >2500 UMIs, >1000 genes detected, less than 10% mitochondrial reads and a novelty >0.8 (log10 of gene number divided by log10 of UMIs). Cells designates as “multiplets” or “undetermined” during demultiplexing were also removed. Data normalization, marker gene expression analyses, clustering, dimensional reduction (UMAP based on top 40 principal components) and differential gene expression analyses were performed with the use of the Seurat package (version 5.0.1)^62^ in R (version 4.3.2). The R package ‘fgsea’ was used for gene set enrichment analyses. Differential gene expression was determined using the ‘FindMarkers’ function in Seurat with default parameters.

### Data analysis and statistics

Statistical testing, including paired and non-paired parametric T-tests , non-parametric Mann-Whitney U-tests, one-way ANOVA with multiple testing adjustment and linear regression analyses were carried out using GraphPad Prism version 8.4.3.

## Acknowledgement

We thank Arif Ekici for performing the scRNAseq runs and providing the raw data files, Jochen Mattner and Stefan Wirtz for providing the Tie2^Cre^_Arg1^fl/fl^ and Tie2^Cre^_RORα^fl/fl^ lines, respectively, Daniela Döhler and Kirstin Castiglione for excellent technical support and members of the Voehringer lab for helpful discussions. This work was supported in part by grants from the Deutsche Forschungsgemeinschaft (RTG2740 project A7A (447268119), VO944/13-1 (500725705), and VO944/14-1 (551748372) as part of project 18 within the Priority Program SPP2332 “Physics of Parasitism”)

## Data Availability Statement

All primary data are available upon reasonable request to the corresponding author

## References

1. Loukas, A., Hotez, P.J., Diemert, D., Yazdanbakhsh, M., McCarthy, J.S., Correa-Oliveira, R., Croese, J., and Bethony, J.M. (2016). Hookworm infection. Nature Reviews Disease Primers 2. 10.1038/nrdp.2016.88.

2. Jõgi, N.O., Kitaba, N., Storaas, T., Schlünssen, V., Triebner, K., Holloway, J.W., Horsnell, W.G.C., Svanes, C., and Bertelsen, R.J. (2022). Ascaris exposure and its association with lung function, asthma, and DNA methylation in Northern Europe. Journal of Allergy and Clinical Immunology, 1-10. 10.1016/j.jaci.2021.11.013.

3. Fan, E.K.Y., and Fan, J. (2018). Regulation of alveolar macrophage death in acute lung inflammation. Respiratory Research 19, 50. 10.1186/s12931-018-0756-5.

4. Marsland, B.J., Kurrer, M., Reissmann, R., Harris, N.L., and Kopf, M. (2008). Nippostrongylus brasiliensis infection leads to the development of emphysema associated with the induction of alternatively activated macrophages. European Journal of Immunology 38, 479–488. 10.1002/eji.200737827.

5. Mearns, H., Horsnell, W.G.C., Hoving, J.C., Dewals, B., Cutler, A.J., Kirstein, F., Myburgh, E., Arendse, B., and Brombacher, F. (2008). Interleukin-4-promoted T helper 2 responses enhance Nippostrongylus brasiliensis-induced pulmonary pathology. Infection and Immunity 76, 5535–5542. 10.1128/IAI.00210-08.

6. Horsnell, W.G.C., Vira, A., Kirstein, F., Mearns, H., Hoving, J.C., Cutler, A.J., Dewals, B., Myburgh, E., Kimberg, M., Arendse, B., et al. (2011). IL-4Rα-responsive smooth muscle cells contribute to initiation of TH2 immunity and pulmonary pathology in Nippostrongylus brasiliensis infections. Mucosal Immunology 4, 83–92. 10.1038/mi.2010.46.

7. Sutherland, T.E., Logan, N., Rückerl, D., Humbles, A.A., Allan, S.M., Papayannopoulos, V., Stockinger, B., Maizels, R.M., and Allen, J.E. (2014). Chitinase-like proteins promote IL-17-mediated neutrophilia in a tradeoff between nematode killing and host damage. Nature Immunology 15, 1116–1125. 10.1038/ni.3023.

8. Ajendra, J., Chenery, A.L., Parkinson, J.E., Chan, B.H.K., Pearson, S., Colombo, S.A.P., Boon, L., Grencis, R.K., Sutherland, T.E., and Allen, J.E. (2020). IL-17A both initiates, via IFNγ suppression, and limits the pulmonary type-2 immune response to nematode infection. Mucosal Immunology 13, 958–968. 10.1038/s41385-020-0318-2.

9. Bouchery, T., Kyle, R., Shepherd, A., Filbey, K., Smith, A., Harvie, M., Painter, G., Johnston, K., Ferguson, P., Jain, R., et al. (2015). ILC2s and T cells cooperate to ensure maintenance of M2 macrophages for lung immunity against hookworms. Nature Communications. 10.1038/ncomms7970.

10. Varela, F., Symowski, C., Pollock, J., Wirtz, S., and Voehringer, D. (2022). IL-4/IL-13-producing ILC2s are required for timely control of intestinal helminth infection in mice. European Journal of Immunology 52, 1925–1933. 10.1002/eji.202249892.

11. Jarick, K.J., Topczewska, P.M., Jakob, M.O., Yano, H., Arifuzzaman, M., Gao, X., Boulekou, S., Stokic-Trtica, V., Leclère, P.S., Preußer, A., et al. (2022). Non-redundant functions of group 2 innate lymphoid cells. Nature. 10.1038/s41586-022-05395-5.

12. Thawer, S.G., Horsnell, W.G.C., Darby, M., Hoving, J.C., Dewals, B., Cutler, A.J., Lang, D., and Brombacher, F. (2014). Lung-resident CD4+ T cells are sufficient for IL-4R-dependent recall immunity to Nippostrongylus brasiliensis infection. Mucosal Immunology 7, 239–248. 10.1038/mi.2013.40.

13. Roberts, J., Chevalier, A., Hawerkamp, H.C., Yeow, A., Matarazzo, L., Schwartz, C., Hams, E., and Fallon, P.G. (2023). Retinoic Acid–Related Orphan Receptor α Is Required for Generation of Th2 Cells in Type 2 Pulmonary Inflammation. The Journal of Immunology. 10.4049/jimmunol.2200896.

14. Chen, F., Wu, W., Millman, A., Craft, J.F., Chen, E., Patel, N., Boucher, J.L., Urban, J.F., Kim, C.C., and Gause, W.C. (2014). Neutrophils prime a long-lived effector macrophage phenotype that mediates accelerated helminth expulsion. Nature Immunology 15, 938–946. 10.1038/ni.2984.

15. Giacomin, P.R., Gordon, D.L., Botto, M., Daha, M.R., Sanderson, S.D., Taylor, S.M., and Dent, L.A. (2008). The role of complement in innate, adaptive and eosinophil-dependent immunity to the nematode Nippostrongylus brasiliensis. Molecular Immunology 45, 446–455. 10.1016/j.molimm.2007.05.029.

16. Chen, F., El-Naccache, D.W., Ponessa, J.J., Lemenze, A., Espinosa, V., Wu, W., Lothstein, K., Jin, L., Antao, O., Weinstein, J.S., et al. (2022). Helminth resistance is mediated by differential activation of recruited monocyte-derived alveolar macrophages and arginine depletion. Cell Reports 38, 110215–110215. 10.1016/j.celrep.2021.110215.

17. Exposure to lung-migrating helminth protects against murine SARS-CoV-2 infection through macrophage-dependent T cell activation. Science immunology. 10.1126/sciimmunol.adf8161.

18. Coakley, G., and Harris, N.L. (2020). Interactions between macrophages and helminths 10.1111/pim.12717.

19. Guilliams, M., De Kleer, I., Henri, S., Post, S., Vanhoutte, L., De Prijck, S., Deswarte, K., Malissen, B., Hammad, H., and Lambrecht, B.N. (2013). Alveolar macrophages develop from fetal monocytes that differentiate into long-lived cells in the first week of life via GM-CSF. Journal of Experimental Medicine 210, 1977–1992. 10.1084/jem.20131199.

20. Tan, S.Y.S., and Krasnow, M.A. (2016). Developmental origin of lung macrophage diversity. Development 143, 1318–1327. 10.1242/dev.129122.

21. Saluzzo, S., Gorki, A.D., Rana, B.M.J., Martins, R., Scanlon, S., Starkl, P., Lakovits, K., Hladik, A., Korosec, A., Sharif, O., et al. (2017). First-Breath-Induced Type 2 Pathways Shape the Lung Immune Environment. Cell Reports 18, 1893–1905. 10.1016/j.celrep.2017.01.071.

22. Westphalen, K., Gusarova, G.A., Islam, M.N., Subramanian, M., Cohen, T.S., Prince, A.S., and Bhattacharya, J. (2014). Sessile alveolar macrophages communicate with alveolar epithelium to modulate immunity. Nature 506, 503–506. 10.1038/nature12902.

23. Neupane, A.S., Willson, M., Chojnacki, A.K., Vargas E Silva Castanheira, F., Morehouse, C., Carestia, A., Keller, A.E., Peiseler, M., DiGiandomenico, A., Kelly, M.M., et al. (2020). Patrolling Alveolar Macrophages Conceal Bacteria from the Immune System to Maintain Homeostasis. Cell 183, 110–125.e111. 10.1016/j.cell.2020.08.020.

24. Chakarov, S., Lim, H.Y., Tan, L., Lim, S.Y., See, P., Lum, J., Zhang, X.-M., Foo, S., Nakamizo, S., Duan, K., et al. (2019). Two distinct interstitial macrophage populations coexist across tissues in specific subtissular niches. Science 363, eaau0964. 10.1126/science.aau0964.

25. Ural, B.B., Yeung, S.T., Damani-Yokota, P., Devlin, J.C., de Vries, M., Vera-Licona, P., Samji, T., Sawai, C.M., Jang, G., Perez, O.A., et al. (2020). Identification of a nerve-associated, lung-resident interstitial macrophage subset with distinct localization and immunoregulatory properties. Science Immunology 5. 10.1126/sciimmunol.aax8756.

26. Aegerter, H., Kulikauskaite, J., Crotta, S., Patel, H., Kelly, G., Hessel, E.M., Mack, M., Beinke, S., and Wack, A. (2020). Influenza-induced monocyte-derived alveolar macrophages confer prolonged antibacterial protection. Nature Immunology 21, 145–157. 10.1038/s41590-019-0568-x.

27. Feo-Lucas, L., Godio, C., Minguito de la Escalera, M., Alvarez-Ladrón, N., Villarrubia, L.H., Vega-Pérez, A., González-Cintado, L., Domínguez-Andrés, J., García-Fojeda, B., Montero-Fernández, C., et al. (2023). Airway allergy causes alveolar macrophage death, profound alveolar disorganization and surfactant dysfunction. Front Immunol 14, 1125984. 10.3389/fimmu.2023.1125984.

28. Kitur, K., Parker, D., Nieto, P., Ahn, D.S., Cohen, T.S., Chung, S., Wachtel, S., Bueno, S., and Prince, A. (2015). Toxin-Induced Necroptosis Is a Major Mechanism of Staphylococcus aureus Lung Damage. PLOS Pathogens 11, e1004820. 10.1371/journal.ppat.1004820.

29. Aegerter, H., Lambrecht, B.N., and Jakubzick, C.V. (2022). Biology of lung macrophages in health and disease. Immunity 55, 1564–1580. 10.1016/j.immuni.2022.08.010.

30. Rodriguez-Rodriguez, L., Gillet, L., and Machiels, B. (2023). Shaping of the alveolar landscape by respiratory infections and long-term consequences for lung immunity. Front Immunol 14, 1149015. 10.3389/fimmu.2023.1149015.

31. Finlay, C.M., Parkinson, J.E., Zhang, L., Chan, B.H.K., Ajendra, J., Chenery, A., Morrison, A., Kaymak, I., Houlder, E.L., Murtuza Baker, S., et al. (2023). T helper 2 cells control monocyte to tissue-resident macrophage differentiation during nematode infection of the pleural cavity. Immunity 56, 1064–1081.e1010. 10.1016/j.immuni.2023.02.016.

32. Gibbons, M.A., MacKinnon, A.C., Ramachandran, P., Dhaliwal, K., Duffin, R., Phythian-Adams, A.T., van Rooijen, N., Haslett, C., Howie, S.E., Simpson, A.J., et al. (2011). Ly6Chi Monocytes Direct Alternatively Activated Profibrotic Macrophage Regulation of Lung Fibrosis. American Journal of Respiratory and Critical Care Medicine 184, 569–581. 10.1164/rccm.201010-1719OC.

33. T’Jonck, W., and Bain, C.C. (2023). The role of monocyte-derived macrophages in the lung: It’s all about context. The International Journal of Biochemistry & Cell Biology 159, 106421. 10.1016/j.biocel.2023.106421.

34. Hilligan, K.L., Oyesola, O.O., Namasivayam, S., Howard, N., Clancy, C.S., Oland, S.D., Garza, N.L., Lafont, B.A.P., Johnson, R.F., Mayer-Barber, K.D., et al. (2022). Helminth exposure protects against murine SARS-CoV-2 infection through macrophage dependent T cell activation. bioRxiv : the preprint server for biology. 10.1101/2022.11.09.515832.

35. Influenza-trained mucosal-resident alveolar macrophages confer long-term antitumor immunity in the lungs. Nature Immunology. 10.1038/s41590-023-01428-x.

36. Dietschmann, A., Ruhl, A., Murray, P.J., Günther, C., Becker, C., Fallon, P., and Voehringer, D. (2023). Th2-dependent disappearance and phenotypic conversion of mouse alveolar macrophages. European Journal of Immunology 53, 2350475. 10.1002/eji.202350475.

37. Symowski, C., and Voehringer, D. (2019). Th2 cell-derived IL-4/IL-13 promote ILC2 accumulation in the lung by ILC2-intrinsic STAT6 signaling in mice. European Journal of Immunology 49, 1421–1432. 10.1002/eji.201948161.

38. Lucas, D.D., and Ens, M. Holding glycolysis in check though Alox15 activity is required for macrophage M2 1 commitment and function in tissue repair and anti-helminth immunity. 10.1101/2024.03.26.586755.

39. Harvie, M., Camberis, M., Tang, S.C., Delahunt, B., Paul, W., and Le Gros, G. (2010). The lung is an important site for priming CD4 T-cell-mediated protective immunity against gastrointestinal helminth parasites. Infection and Immunity 78, 3753–3762. 10.1128/IAI.00502-09.

40. Knipfer, L., Schulz-Kuhnt, A., Kindermann, M., Greif, V., Symowski, C., Voehringer, D., Neurath, M.F., Atreya, I., and Wirtz, S. (2019). A CCL1/CCR8-dependent feed-forward mechanism drives ILC2 functions in type 2-mediated inflammation. J Exp Med 216, 2763–2777. 10.1084/jem.20182111.

41. Harvie, M., Camberis, M., and Gros, G.L. (2013). Development of CD4 T cell dependent immunity against N. brasiliensis infection. Frontiers in Immunology 4, 1–5. 10.3389/fimmu.2013.00074.

42. Chen, F., Liu, Z., Wu, W., Rozo, C., Bowdridge, S., Rooijen, N.V., Jr, J.F.U., Wynn, T.A., and William, C. (2012). An essential role for the Th2-type response in limiting tissue damage during helminth infection. 18, 260–266. 10.1038/nm.2628.An.

43. Lidia, B., Grace, C.Y., Mar, C.-C., Andrea, T., D, H.L., Yong, K., S, W.J., Paula, L.-L., T, S.E., Facundo, P., et al. (2017). Macrophage function in tissue repair and remodeling requires IL-4 or IL-13 with apoptotic cells. Science 356, 1072–1076. 10.1126/science.aai8132.

44. Obata-Ninomiya, K., Ishiwata, K., Tsutsui, H., Nei, Y., Yoshikawa, S., Kawano, Y., Minegishi, Y., Ohta, N., Watanabe, N., Kanuka, H., and Karasuyama, H. (2013). The skin is an important bulwark of acquired immunity against intestinal helminths. Journal of Experimental Medicine 210, 2583–2595. 10.1084/jem.20130761.

45. Westermann, S., Schubart, C., Dietschmann, A., Castiglione, K., Radtke, D., and Voehringer, D. (2023). Th2-dependent STAT6-regulated genes in intestinal epithelial cells mediate larval trapping during secondary Heligmosomoides polygyrus bakeri infection. PLoS Pathog 19, e1011296. 10.1371/journal.ppat.1011296.

46. Chetty, A., Darby, M.G., Pillaye, J., Taliep, A., Cunningham, A.F., O’Shea, M.K., Katawa, G., Layland, L.E., Ritter, M., and Horsnell, W.G.C. (2023). Induction of Siglec-F(hi)CD101(hi) eosinophils in the lungs following murine hookworm Nippostrongylus brasiliensis infection. Front Immunol 14, 1170807. 10.3389/fimmu.2023.1170807.

47. Cui, A., Huang, T., Li, S., Ma, A., Pérez, J.L., Sander, C., Keskin, D.B., Wu, C.J., Fraenkel, E., and Hacohen, N. (2024). Dictionary of immune responses to cytokines at single-cell resolution. Nature 625, 377–384. 10.1038/s41586-023-06816-9.

48. Arguello, R.J., Combes, A.J., Char, R., Gigan, J.P., Baaziz, A.I., Bousiquot, E., Camosseto, V., Samad, B., Tsui, J., Yan, P., et al. (2020). SCENITH: A Flow Cytometry-Based Method to Functionally Profile Energy Metabolism with Single-Cell Resolution. Cell Metab 32, 1063–1075 e1067. 10.1016/j.cmet.2020.11.007.

49. Heieis, G.A., Patente, T.A., Almeida, L., Vrieling, F., Tak, T., Perona-Wright, G., Maizels, R.M., Stienstra, R., and Everts, B. (2023). Metabolic heterogeneity of tissue-resident macrophages in homeostasis and during helminth infection. Nat Commun 14, 5627. 10.1038/s41467-023-41353-z.

50. Heieis, G.A., Patente, T.A., Almeida, L., Vrieling, F., Tak, T., Perona-Wright, G., Maizels, R.M., Stienstra, R., and Everts, B. (2023). Metabolic heterogeneity of tissue-resident macrophages in homeostasis and during helminth infection. Nature Communications 14. 10.1038/s41467-023-41353-z.

51. Rai, G., Joshi, N., Perry, S., Yasgar, A., Schultz, L., Jung, J.E., Liu, Y., Terasaki, Y., Diaz, G., Kenyon, V., et al. (2010). Discovery of ML351, a Potent and Selective Inhibitor of Human 15-Lipoxygenase-1. In Probe Reports from the NIH Molecular Libraries Program.

52. Barth, M.W., Hendrzak, J.A., Melnicoff, M.J., and Morahan, P.S. (1995). Review of the macrophage disappearance reaction. Journal of Leukocyte Biology 57, 361–367. 10.1002/jlb.57.3.361.

53. Shibata, S., Miyake, K., Tateishi, T., Yoshikawa, S., Yamanishi, Y., Miyazaki, Y., Inase, N., and Karasuyama, H. (2018). Basophils trigger emphysema development in a murine model of COPD through IL-4–mediated generation of MMP-12–producing macrophages. Proceedings of the National Academy of Sciences 115, 13057–13062. 10.1073/pnas.1813927115.

54. Haider, P., Kral-Pointner, J.B., Salzmann, M., Moik, F., Bleichert, S., Schrottmaier, W.C., Kaun, C., Brekalo, M., Fischer, M.B., Speidl, W.S., et al. (2022). Interleukin-4 receptor alpha signaling regulates monocyte homeostasis. The FASEB Journal 36, e22532. 10.1096/fj.202101672RR.

55. Guilliams, M., De Kleer, I., Henri, S., Post, S., Vanhoutte, L., De Prijck, S., Deswarte, K., Malissen, B., Hammad, H., and Lambrecht, B.N. (2013). Alveolar macrophages develop from fetal monocytes that differentiate into long-lived cells in the first week of life via GM-CSF. J Exp Med 210, 1977–1992. 10.1084/jem.20131199.

56. Svedberg, F.R., Brown, S.L., Krauss, M.Z., Campbell, L., Sharpe, C., Clausen, M., Howell, G.J., Clark, H., Madsen, J., Evans, C.M., et al. (2019). The lung environment controls alveolar macrophage metabolism and responsiveness in type 2 inflammation. Nature Immunology 20, 571–580. 10.1038/s41590-019-0352-y.

57. Subramanian, S., Busch, C.J.-L., Molawi, K., Geirsdottir, L., Maurizio, J., Vargas Aguilar, S., Belahbib, H., Gimenez, G., Yuda, R.A.A., Burkon, M., et al. (2022). Long-term culture-expanded alveolar macrophages restore their full epigenetic identity after transfer in vivo. Nature Immunology 23, 458–468. 10.1038/s41590-022-01146-w.

58. Li, F., Piattini, F., Pohlmeier, L., Feng, Q., Rehrauer, H., and Kopf, M. (2022). Monocyte-derived alveolar macrophages autonomously determine severe outcome of respiratory viral infection. Science Immunology 7, eabj5761. 10.1126/sciimmunol.abj5761.

59. Iliakis, C.S., Kulikauskaite, J., Aegerter, H., Li, F., Piattini, F., Jakubzick, C.V., Guilliams, M., Kopf, M., and Wack, A. (2023). The role of recruitment versus training in influenza-induced lasting changes to alveolar macrophage function. Nature Immunology 24, 1639–1641. 10.1038/s41590-023-01602-1.

60. Heieis, G.A., Patente, T.A., Almeida, L., Vrieling, F., Tak, T., Perona-Wright, G., Maizels, R.M., Stienstra, R., and Everts, B. (2023). Metabolic heterogeneity of tissue-resident macrophages in homeostasis and during helminth infection. Nature Communications 14, 5627. 10.1038/s41467-023-41353-z.

61. Snodgrass, R.G., and Brüne, B. (2019). Regulation and Functions of 15-Lipoxygenases in Human Macrophages. Front Pharmacol 10, 719. 10.3389/fphar.2019.00719.

62. Pernet, E., Sun, S., Sarden, N., Gona, S., Nguyen, A., Khan, N., Mawhinney, M., Tran, K.A., Chronopoulos, J., Amberkar, D., et al. (2023). Neonatal imprinting of alveolar macrophages via neutrophil-derived 12-HETE. Nature 614, 530–538. 10.1038/s41586-022-05660-7.

63. Huang, J.T., Welch, J.S., Ricote, M., Binder, C.J., Willson, T.M., Kelly, C., Witztum, J.L., Funk, C.D., Conrad, D., and Glass, C.K. (1999). Interleukin-4-dependent production of PPAR-gamma ligands in macrophages by 12/15-lipoxygenase. Nature 400, 378–382. 10.1038/22572.

64. Abrial, C., Grassin-Delyle, S., Salvator, H., Brollo, M., Naline, E., and Devillier, P. (2015). 15-Lipoxygenases regulate the production of chemokines in human lung macrophages. British Journal of Pharmacology 172, 4319–4330. 10.1111/bph.13210.

65. Minutti, C.M., Jackson-Jones, L.H., García-Fojeda, B., Knipper, J.A., Sutherland, T.E., Logan, N., Rinqvist, E., Guillamat-Prats, R., Ferenbach, D.A., Artigas, A., et al. (2017). Local amplifiers of IL-4Ra-mediated macrophage activation promote repair in lung and liver. Science 356, 1076–1080. 10.1126/science.aaj2067.

66. DeBerge, M., Glinton, K., Lantz, C., Ge, Z.-D., Sullivan, D.P., Patil, S., Lee, B.R., Thorp, M.I., Mullick, A., Yeh, S., et al. (2025). Mechanical regulation of macrophage metabolism by allograft inflammatory factor 1 leads to adverse remodeling after cardiac injury. Nature Cardiovascular Research 4, 83–101. 10.1038/s44161-024-00585-y.

67. Wei, M., Zhou, G., Chen, L., Zhang, Y., Ma, W., Gao, L., and Gao, G. (2024). The prognostic and immune significance of PLBD1 in pan-cancer and its roles in proliferation and invasion of glioma. J Cancer 15, 3857–3872. 10.7150/jca.96365.

68. Voehringer, D., van Rooijen, N., and Locksley, R.M. (2007). Eosinophils develop in distinct stages and are recruited to peripheral sites by alternatively activated macrophages. J Leukoc Biol 81, 1434–1444. 10.1189/jlb.1106686.

69. Knipfer, L., Schulz-Kuhnt, A., Kindermann, M., Greif, V., Symowski, C., Voehringer, D., Neurath, M.F., Atreya, I., and Wirtz, S. (2019). A CCL1/CCR8-dependent feed-forward mechanism drives ILC2 functions in type 2-mediated inflammation. Journal of Experimental Medicine 216, 2763–2777. 10.1084/jem.20182111.

70. Oeser, K., Schwartz, C., and Voehringer, D. (2015). Conditional IL-4/IL-13-deficient mice reveal a critical role of innate immune cells for protective immunity against gastrointestinal helminths. Mucosal Immunology 8, 672–682. 10.1038/mi.2014.101.

71. Herndler-Brandstetter, D., Ishigame, H., Shinnakasu, R., Plajer, V., Stecher, C., Zhao, J., Lietzenmayer, M., Kroehling, L., Takumi, A., Kometani, K., et al. (2018). KLRG1+ Effector CD8+ T Cells Lose KLRG1, Differentiate into All Memory T Cell Lineages, and Convey Enhanced Protective Immunity. Immunity 48, 716–729.e718. 10.1016/j.immuni.2018.03.015.

72. Haag, S.M., and Murthy, A. (2021). Murine Monocyte and Macrophage Culture. Bio Protoc 11, e3928. 10.21769/BioProtoc.3928.

