## Supplementary Figures and Table for "ILC2s govern imprinting of alveolar macrophage-mediated immunity upon secondary helminth infection in the lung"

Supplementary Figure 1

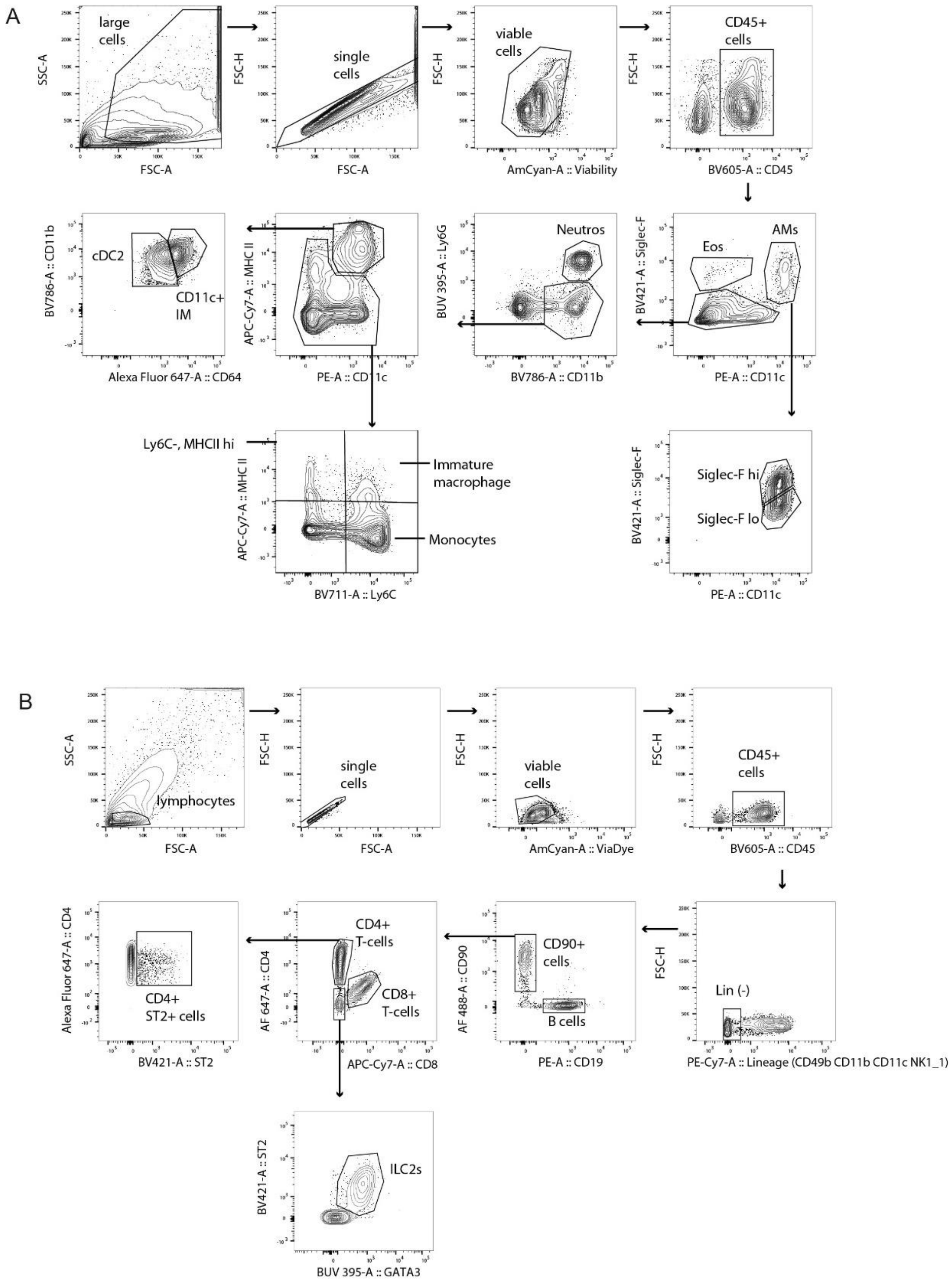


**Supplementary Figure 1. Gating strategies used for analysis of myeloid and lymphoid subsets in the lungs of mice during 2° Nb infection.** A) Gating strategy of myeloid subsets. B) Gating strategy of lymphoid subsets

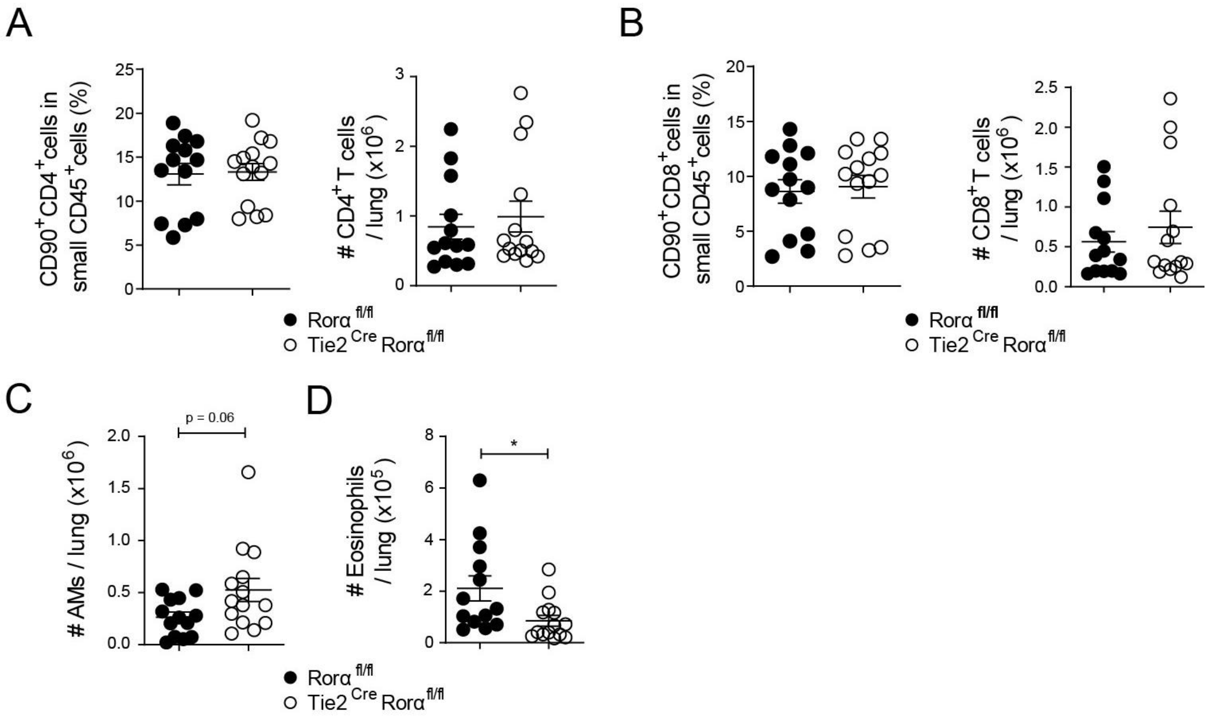


**Supplementary Figure 2. Lung myeloid and lymphoid subsets in the lungs of Tie2$^{Cre}$Rorα$^{fl/fl}$ and littermate control mice during 2° Nb infection.** All data are from lung on day 2 post 2° infection. A) Frequencies and total numbers of CD4$^{+}$ T cells. B) Frequencies and total numbers of CD8$^{+}$ T cells. C) Total number of alveolar macrophages (AMs). D) Total number of eosinophils. Data are pooled from 4 independent experiments (n = 3 -4 mice per group). Mann-Whitney U-tests used for statistical analysis. Data include the mean ± SEM. *, $p < 0.05$.

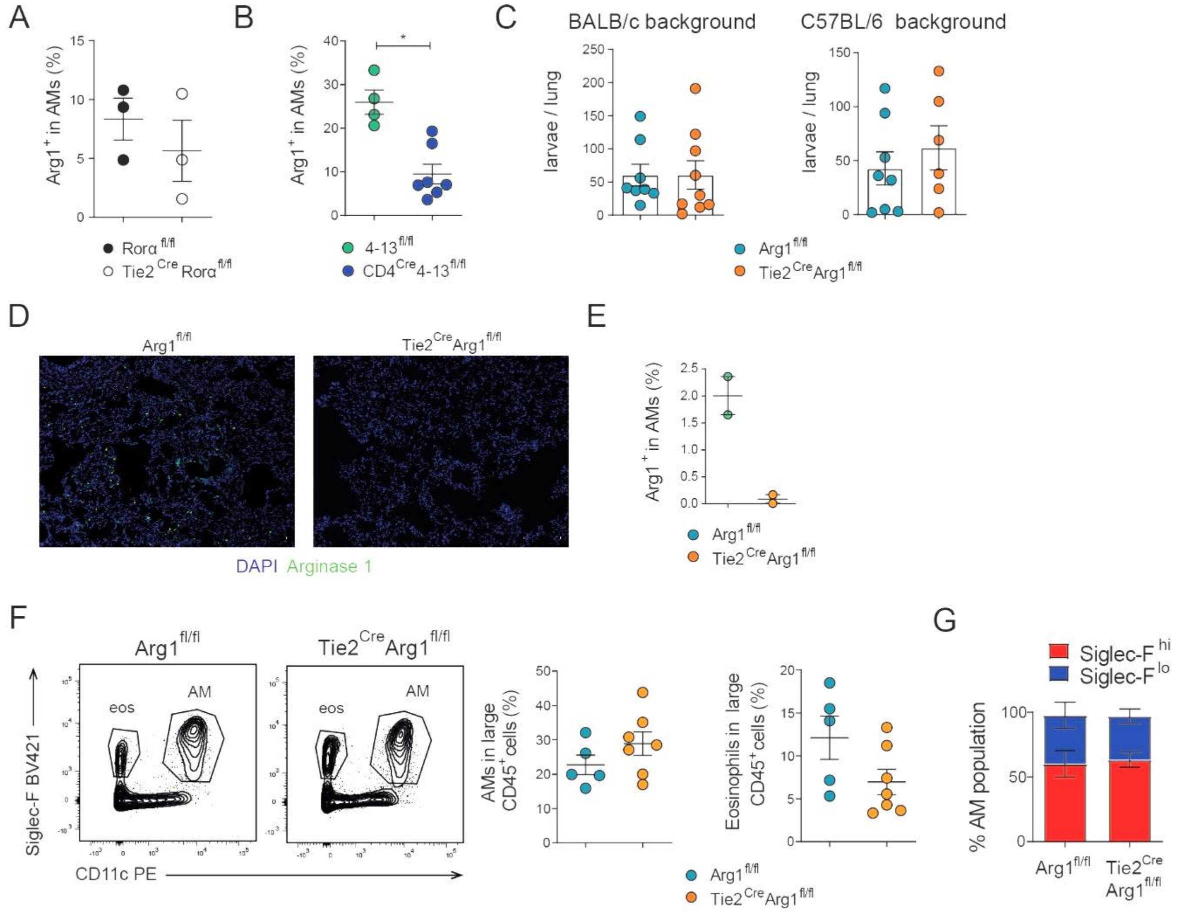


**Supplementary Figure 3. Role of Arginase 1 during 2° Nb infection.** All data are from lungs of mice analysed at day 2 post 2° infection with Nb. A and B) Frequency of Arg1$^{+}$ cells within the alveolar macrophage (AM) population of indicated mice. C) Number of worms in the lungs of Tie2$^{Cre}$Arg1$^{fl/fl}$ and control mice from two different mouse backgrounds at day 2 post 2° infection. D) Immunofluorescent staining of Arginase 1 in cryosections of lung tissue from Tie2$^{Cre}$ Arg1$^{fl/fl}$ and control mice. E) Intracellular staining for Arginase 1 in AM of indicated mice. F) Representative flow cytometry plots showing relative levels of AMs and eosinophils in indicated mice. G) Relative proportions of Siglec-F$^{hi}$ and Siglec-F$^{lo}$ AMs in indicated mice. Data in A, B, D and E from one experiment, representative of two experiments. Data in C pooled from 2-3 independent experiments (n = 2-3 mice per group). Data in F and G pooled from three independent experiments (n = 1-3 mice per group). Unpaired T-tests were used for statistical anaylsis. Data include the mean ± SEM. *, $p < 0.05$.

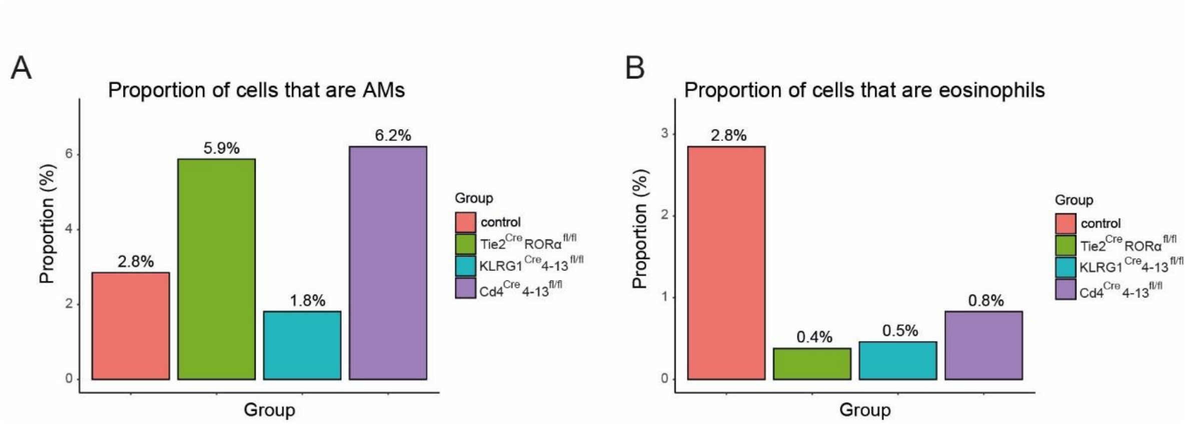


**Supplementary Figure 4. Impact of ILC2s and Th2 cells on AMs and eosinophils.** Tie2$^{Cre}$Rorα$^{fl/fl}$, KLRG1$^{Cre}$4-13$^{fl/fl}$ , Cd4$^{Cre}$4-13$^{fl/fl}$ mice and littermate controls were infected with Nb and 9 days later lung single cell suspensions were subjected to scRNA-seq analysis. Cell-cluster identities are as in Figure 4. A) Proportion of all cells (after filtering) belonging to cluster 8 (comprising AMs and AM-like cells) in respective groups. B) Proportion of all cells belonging to cluster 14 (eosinophils) in respective groups. Data in A and B are from one single-cell RNAseq experiment (n = 2 mice per group).

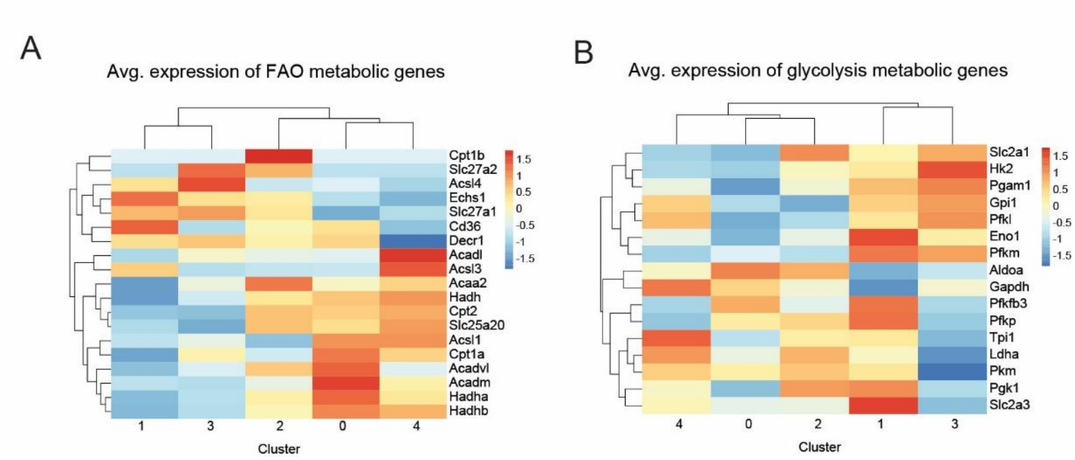


**Supplementary Figure 5. Analyses of expression of select genes involved in fatty acid oxidation and glycolysis. A)** Heatmap showing relative expression of fatty acid oxidation genes in respective macrophage subclusters from Figure 4B. B) Heatmap showing relative expression of glycolysis genes in respective macrophage subclusters from Figure 4B.

### Supplementary Table 1. List of reagents, consumables and resources

| Reagent or Resource | Source | Identifier |
|---|---|---|
| **Antibodies: Flow cytometry** | | |
| Anti-mouse CD3– PE-Cy7 (17A2) | BioLegend | 100220 |
| Anti-mouse CD4 - APC (Gk1.5) | BioLegend | 100412 |
| Anti-Mouse CD8– APC-Cy7 (53-6.7) | eBioscience | 47-0081-82 |
| Anti-mouse CD11b – BV786 /PE-Cy7 (M1/70) | BioLegend | 101216 |
| Anti-mouse CD11c – PE/ PE-Cy7 / BUV737 (N418) | eBioscience | PE 12-0114-82<br>PE-Cy7 25-0114-82<br>BUV737 367-0114-82 |
| Anti-mouse CD19 – PE (eBio 8D9) | eBioscience | 12-0199-42 |
| Anti-mouse CD45 – BV605 (30-F11) | BioLegend | 103139 |
| Anti-mouse CD49b (DX5) | eBioscience | 14-5971-85 |
| Anti-mouse CD64 – APC (X54-5/7.1) | BioLegend | 139306 |
| Anti-mouse CD90 – FITC (30-H12) | eBioscience | 11-0903-82 |
| Anti-mouse CD127 – BUV786 (A7R34) | BioLegend | 135037 |
| Anti-mouse Arginase 1 – AF488 (Alex-F5) | Thermo Fischer Scientific | 53-3697-82 |
| Anti-mouse GATA3 – (L50-823) | Becton Dickinson | 558686 |
| Anti-mouse KLRG1- BV711 (2F1) | Becton Dickinson | 564014 |
| Anti-mouse Ly6C – BV711 (AL-21) | Becton Dickinson | 755195 |
| Anti-mouse Ly6G – APC-Cy7 (1A8) | BioLegend | 127624 |
| Anti-mouse MerTk – PE-Cy7 (DS5MMER) | eBioscience | 25-5751-82 |
| Anti-mouse MHC II – BUV395 (2G9 ) | Becton Dickinson | 569244 |
| Anti-mouse Nk1.1 – PE-Cy7 (PK136) | eBiosciences | 25-5941-82 |
| Anti- puromycin – AF647 (12D10) | Merck | |
| Anti-puromycin – AF488 (2A4) | BioLegend | 381505 |
| Anti-mouse PD-L-2 – PE-Cy7 (TY25) | BioLegend | 107214 |
| Anti-mouse ST2- BV421 (U29-93) | Becton Dickinson | 566310 |
| Anti-mouse SiglecF – BV421 (E50-2440) | Becton Dickinson | 562681 |
| Fixable Viability Dye eFluor™ 506 | eBioscience | 65-0865-18 |
| **Antibodies: Immunofluorescence** | | |
| Anti-mouse Relm-α (polyclonal) | Abcam | ab272692 |
| Anti-mouse Arginase-1 Rabbit mAb (D4E3M) | Cell signalling | 93668S |
| C1qC Polyclonal antibody | Proteintech | 16889-1-AP |
| Anti-mouse CD68 - AF647 / AF488 (FA-11) | BioLegend | 137004 |
| Rat anti - mouse SiglecF (E50-2440) | Becton Dickinson | 552126 |
| Goat anti-rat IgG- AF488 | Jackson ImmunoResearch | 112-545-003 |
| Donkey anti-rabbit IgG- AF647 | Jackson ImmunoResearch | 711-605-152 |
| **Oligonucleotides** | | |
| Gapdh-fw | 5'-CGTCTTCACCACCATGGAGA-3' | |
| Gapdh-rv | 5'-CGGCCATCACGCCACAGTTT-3' | |
| Muc5ac_fw | 5'-GGCTAACCTGTGGGCTCTGTGGTA-3' | |
| Muc5ac_rv | 5'-CACAAGCACGCACATAGGAGGACAG-3' | |
| Clca1_fw | 5'-ACCTTCAAAAACGCTGATGTCCTT-3' | |
| Clca1_rv | 5'-ACTTTCCGTTAGTGATACAACTGCC-3' | |

| | | |
|---|---|---|
| Sftpd_fw | 5’-CCAACACCTGCACCCTAGTCATGT-3’ | |
| Sftpd_fw | 5’-GGAGCACCTACTTCTCCTTTGGGC-3’ | |
| Alox15_fw | 5’-CCAAGATGGCTGAGCGGGTTC-3’ | |
| Alox15_rv | 5’-GGCAGTTCGAGCTGGATGGC-3’ | |
| Alox5_fw | 5’-CCCATTGCCATCCAGCTCAACC-3’ | |
| Alox5_rv | 5’-CCGGTGGCATTGGCCTTGTC-3’ | |
| Ptgs1_fw | 5’-CCACTCCCAGAGTCATGAGTCGAAG-3’ | |
| Ptgs1_rv | 5’-CCAGATCTCAGGGATGGTACAGTTGG-3’ | |
| Ptgs2_fw | 5’-GGTGTGAAGGGAAATAAGGAGCTTCC-3’ | |
| Ptgs2_rv | 5’-CACCTCTCCACCAATGACCTGATATTTC-3’ | |
| **Other reagents** | | |
| Liberase TM | Roche | 5401119001 |
| GlutaMax 100x Supplement | Thermo Fisher Scientific | 35050061 |
| Fetal calf/bovine serum (FCS) | Thermo Fisher Scientific | A5256701 |
| Fluoroshield | Sigma-Aldrich | F6182-20ML |
| Tween 20 | AppliChem | A4974.1000 |
| DNase I | Sigma-Aldrich | DN25-100MG |
| Oligomycin A | MedChemExpress | HY-16589 |
| 2-Deoxy-D-glucose | Sigma-Aldrich | D8375-1G |
| Puromycin | Sigma-Aldrich | P8833-10MG |
| Complete Protease Inhibitor Cocktail | Roche | 4693116001 |
| Entellan | Sigma-Aldrich | 1.079.600.500 |
| ML351 (ALOX15 inhibitor) | Hycultec | HY-111310 |
| 4′,6-Diamidin-2-phenylindol (DAPI) | Sigma-Aldrich | 32670 |
| Recombinant mouse IL-4 | R&D systems | 404-ML-025 |
| Recombinant M-CSF | L929 cell line | |
| **Consumables and kits** | | |
| Tungsten carbide beads 3 mm | Qiagen | 6997 |
| RNeasy Mini Kit | Qiagen | 74104 |
| Micro screw cap tubes | Sarstedt | 72.693.005 |
| RNAprotect Tissue Reagent | Qiagen | 76106 |
| Foxp3/Transcription Factor Staining kit | eBiosciences | 00-5523-00 |
| High-Capacity cDNA Reverse Transcription Kit | Thermo Fisher Scientific | 4368813 |
| Pierce BCA Protein Assay Kit | Thermo Fisher Scientific | 23227 |
| Petri dishes, standard | Saerstadt | 82.1472 |
| Dead Cell Removal Kit | Miltenyi | 130-090-101 |
| **Equipment** | | |
| Cellometer Auto T4 cell counter | Nexcelom Bioscience | |
| Bead Ruptor Elite | Biolab products | |
| **Software** | | |
| FlowJo 10.8 | BD Biosciences | |
| ZEN 3.0 Blue Edition | Carl Zeiss | |
| LEGENDplex analysis software | BioLegend | |
| GraphPad Prism | Dotmatics | |
